# Dynamic colonization of microbes and their functions after fecal microbiota transplantation for inflammatory bowel disease

**DOI:** 10.1101/649384

**Authors:** Nathaniel D. Chu, Jessica W. Crothers, Le T.T. Nguyen, Sean M. Kearney, Mark B. Smith, Zain Kassam, Cheryl Collins, Ramnik Xavier, Peter L. Moses, Eric J. Alm

## Abstract

For fecal microbiota transplantation (FMT) to be successful in complex immune diseases like inflammatory bowel disease (IBD), it is assumed that therapeutic microbes and their beneficial functions and immune interactions must colonize the recipient and persist in sufficient quantity and for a long enough period of time to result in a clinical benefit. But few studies have comprehensively profiled the colonization and persistence of transferred microbes along with the transfer of their microbial and immune functions. Using 16S, metagenomic, and immunoglobulin A (IgA) sequencing, we analyzed hundreds of longitudinal microbiome samples from a randomized controlled trial of 12 patients with ulcerative colitis who received fecal transplant or placebo for 12 weeks. We uncovered a range of competitive dynamics among donor and patient strains, showing that persistence of transferred microbes is far from static. Indeed, one patient experienced dramatic loss of donor bacteria 10 weeks into the trial, coinciding with a bloom of pathogenic bacteria and worsening clinical symptoms. We similarly evaluated transfer of microbial functions, including desired ones like butyrate production and unintended ones like antibiotic resistance. By profiling bacteria coated with IgA, we identified IgA-coated bacteria associated with inflammation, and we found that microbial interactions with the host immune system can be transferred across people. This transfer of immune function is likely critical for gut microbiome therapeutics for immune-related diseases. Our findings elucidate the colonization dynamics of gut microbes as well as their functions in the context of FMT to treat a complex disease—information that may provide a critical foundation for the development of more-targeted therapeutics.

## Introduction

Buoyed by early success in recurrent *Clostridium difficile* infections *(1, 2)*, researchers are exploring whether fecal microbiota transplantation (FMT)—the transfer of entire fecal microbial communities from a healthy donor to a sick patient—can treat other microbiome-associated conditions. One of the most promising candidates is inflammatory bowel disease (IBD), a chronic condition characterized by periods of relapse (i.e., “flares”) and remission, which suggest that longitudinal dynamics are key to understanding and treating the disease *(3)*. When compared with healthy individuals, patients suffering from IBD (ulcerative colitis or Crohn’s disease) have distinct gut microbial communities *(4, 5)*. Thus, it has been hypothesized that manipulation of the gut microbiome and its interactions with the gut immune system might improve patient symptoms. Clinical trials have demonstrated that FMT is moderately effective in patients with ulcerative colitis, but the factors driving patient response or nonresponse remain unknown *(6)*.

It is broadly believed that the therapeutic element of FMT is microbes and their functions *(7, 8)*. Many commensal bacteria are thought to promote gut and immune health, for example by the production of butyrate, which plays metabolic *(9)*, regulatory *(10)*, and immune roles *(11–14)* in supporting the gut epithelium. But not all microbial functions are beneficial. Fecal transplant material is rigorously screened for pathogens and large clinical studies have demonstrated its broad safety *(1)*, but the upsurge of antibiotic resistance has raised concerns that fecal transplants could transfer potentially deleterious microbial functions *(15)*.

In addition to the autonomous functions of the microbes themselves, the microbes’ interactions with the gut immune system may also play key roles in disease progression or treatment. The host immune system interacts with gut bacteria by responding to bacterial metabolites *(12, 14)*, sensing direct contact between the host epithelium and bacteria *(16)*, and coating bacteria with immunoglobulin A (IgA)—the main antibody produced in the gut and other mucosal tissues *(17)*. These interactions play a pivotal role in the formation and maintenance of the host’s immune system *(18, 19)*. Since many microbiome-associated diseases—including IBD—are of immune origin, the immune function of the gut microbiome might be the most directly related to host health.

Despite excitement around applying fecal transplantation to IBD, no studies have comprehensively evaluated (1) which microbes transfer and persist across hosts, (2) the microbial functions that accompany them, and (3) whether immune functions of gut bacteria also transfer from donor to recipient. Previous reports of fecal transplants in IBD patients observed variable colonization by bacterial taxa from donors to recipients but did not categorize the functions and immune interactions that were also transferred *(20–23)*. Furthermore, most of these studies used minimal sampling (e.g., single time points before and after a single fecal transplant) and so could not show how transferred bacteria and functions varied over time. Particularly in the case of chronic, inflammatory diseases like IBD, understanding the longitudinal dynamics of transferred microbes and functions would advance our ability to determine why FMT works for some patients and not others and help pave the way for more targeted therapies.

We comprehensively profiled the colonization dynamics of microbes and functions in a small randomized controlled clinical trial of 12 patients who had mild to moderate ulcerative colitis and were treated with FMT. By bringing together analysis of microbial taxa, strains, functions, and immune interaction in this focused clinical cohort, we sought to deeply understand colonization in a limited number of patients to shed light on these dynamics in the context of a complex disease.

## Results

### Study design

We recruited patients at the University of Vermont Medical Center. We deliberately selected two donors with high stool butyrate content—measured by gas chromatography—as loss of butyrate-producing microbes has been associated with inflammation and IBD *(24)*. After a course of broad-spectrum antibiotics (ciprofloxacin and metronidazole for seven days), patients received colonoscopic delivery (in the cecum and terminal ileum) followed by 12 weeks of daily capsules of either fecal transplant material (from one of our two donors) or placebo (Figure 1a). We delivered 120 ml of FMT material (1 g of donor stool per 2.5 ml of material) during colonoscopy and each capsule contained approximately 0.5 g of donor material. We chose to couple antibiotic pretreatment with two transplant delivery methods to maximize each patient’s exposure to donor material and to increase the likelihood that donor bacteria would successfully colonize their new host. To test whether a previously established microbial community could be invaded by new bacteria from low-dosage capsules, a subset of fecal transplant recipients (*n* = 4) received capsule fecal transplant material from an alternate donor for 4 weeks in the middle of the clinical trial, after which they returned to taking material from their original donor (Figure 1a). We collected by mail near-weekly preserved stool samples from these patients during the trial and at an 18-week follow-up. At four time points, we also collected fresh stool samples during clinical check-ins. We sequenced DNA from these stool samples at the Broad Institute (Cambridge, MA) using 16S rDNA sequencing and shotgun metagenomic sequencing, producing datasets comprising an average of ~250,000 16S rDNA sequences and ~30.5 million metagenomic DNA sequences per sample (Table S1). To identify the abundance of different amplicon sequence variants (ASVs, akin to a bacterial species), we processed the 16S rDNA sequences using QIIME 2 *(25)* and DADA2 *(26)*. To track the abundance of bacterial species, we processed the metagenomic sequences using Metaphlan2 *(27)*.

**Fig 1.**
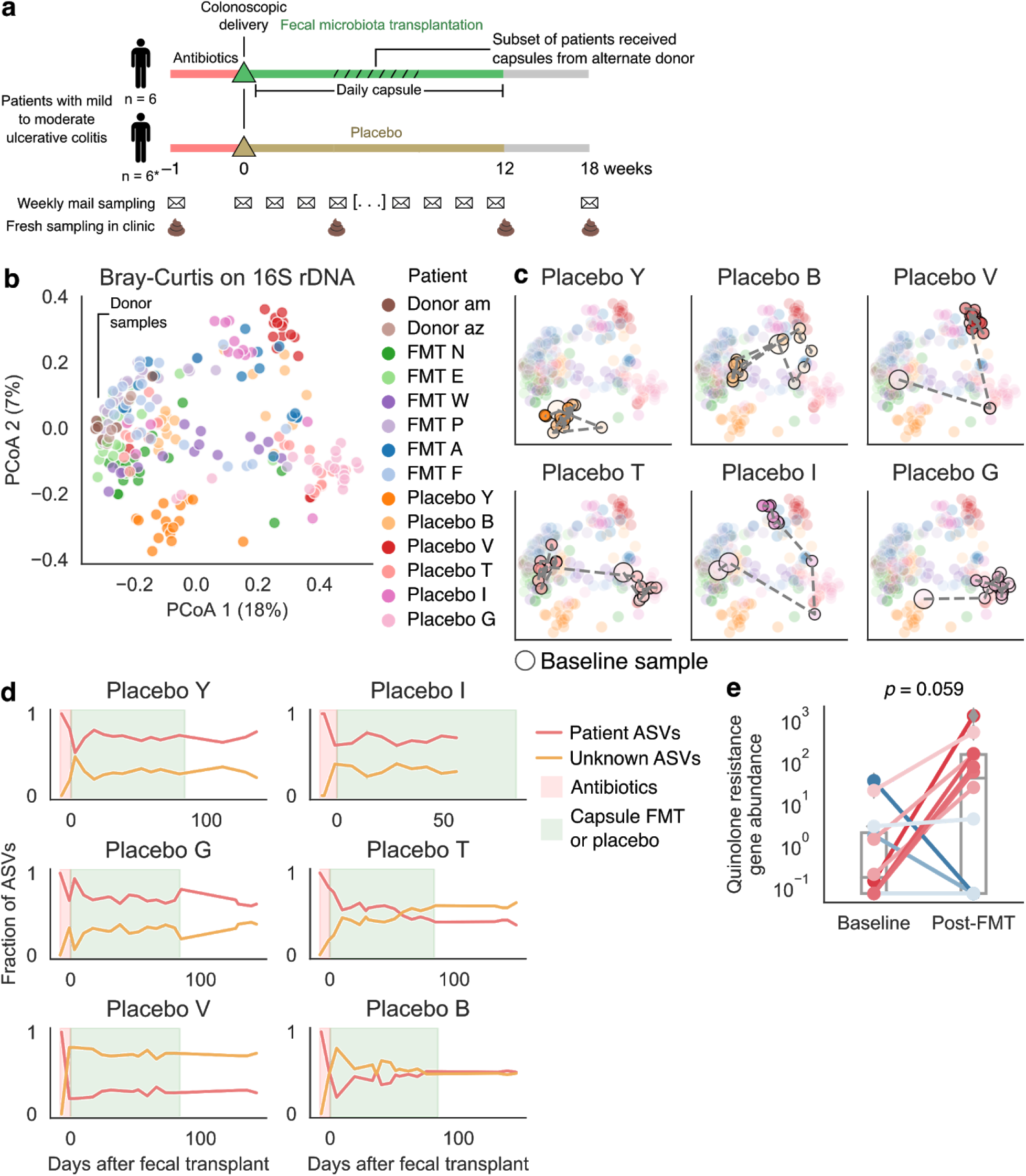
Recovery from antibiotics in IBD patients involved substantial loss and introduction of microbial taxa. (a) Design of the clinical trial and sampling. *One placebo-treated patient had worsening symptoms and dropped out of the trial at 8 weeks. (b) PCoA based on Bray-Curtis distance using 16S. Donor samples clustered on the left-hand side, and patient samples taken during antibiotics treatment tended to cluster toward the right. (c) PCoA trajectories of placebo-treated patients indicated incomplete recovery from antibiotics in most patients. The PCoA space is the same plot as in Figure 1b. The larger circles signify a baseline sample at the beginning of the clinical trial. (d) Tracking microbial sources revealed invasion of many novel bacteria after antibiotics as well as loss of many taxa. Each plot tracks the fraction of ASVs that were identified as coming from the patient (detected in baseline samples) or from an unknown source (bacteria that may have come from the environment or been below our limit of detection in baseline samples). The red region of the figure indicates the course of antibiotics, while the green region indicates the course of capsule therapy (placebo or fecal transplant). (e) Many placebo and transplant recipients exhibited an increase in abundance of quinolone resistance genes in the 10 days after the administration of antibiotics. Lines are colored by the change in resistance. Line color reflects the change, with red lines indicating an increase, and blue lines indicating no change or a decrease. Also see Figure S1.

### Antibiotics destabilized the microbiome of IBD patients

An auxiliary finding that emerged in parallel with our primary results on colonization was that antibiotics destabilized the gut microbial communities of patients receiving placebo, resulting in large-scale microbial gain and loss in these patients. The magnitude of changes we observed was greater than expected in healthy subjects. Previous studies tracking recovery of the gut microbiome in healthy people after broad-spectrum antibiotics observed perturbation followed by consistent recovery of microbial communities—loss of bacteria followed by regain of the same bacteria (Figure S1a) *(28)*.

In contrast, patients in our cohort who received placebo exhibited diverse trajectories in their microbiomes. We calculated average beta diversity between patient samples and donor samples using Bray-Curtis distance and 16S rDNA and metagenomic species datasets. Visualizing the differences between microbial communities using PCoA (Figure 1b, S1b), we found that many placebo-treated patients ended the trial with a microbial community composition very different from where they started (Figure 1c, S1c–d). To see whether placebo-treated patients recovered the same bacteria that they lost while taking antibiotics, we categorized bacteria in each patient’s samples by their putative sources, including those detected in the patient’s baseline samples, taken at the beginning of the clinical trial (“Patient”, orange lines), and those not detected (“Unknown”, yellow lines) (Figure 1d, S1e). This latter category potentially included newly colonized bacteria from the environment, as well as endogenous patient bacteria that were under our detection limit. Most patients receiving placebo regained some endogenous bacteria lost during antibiotics. But, surprisingly, many patients (Placebo B, Placebo V, Placebo T) ended the trial with more bacteria from unknown sources than from themselves, reflecting a remarkable turnover in their gut microbiomes. Although individual patients lost variable sets of taxa, a number of taxa were lost by multiple patients, including butyrate-producing *Subdoligranulum*, *Faecalibacterium*, and *Alistipes*, and commensals *Bacteroides* and *Dorea* (Table S2, S3).

In addition, we found that antibiotics triggered a short-lived increase in specific antibiotic resistance genes across all patients (Methods, Figure 1e). Specifically, the abundance of quinolone resistance genes increased immediately after antibiotics, likely reflecting selection pressure from ciprofloxacin, a quinolone (Figure 1e). These increases were not maintained over time (Figure S1f); neither did we observe temporary or lasting increases in other classes of antibiotic resistance (Figure S1f) or antibiotic resistance overall (Figure S1g,h). Placebo-treated patients also had a greater burden of tetracycline and aminoglycoside resistance genes than did fecal transplant recipients during the clinical trial period, although these levels were not appreciably higher than in baseline samples (Figure S1i,j). Taken together, our results emphasize that disease context can influence the gut microbiome’s ability to recover from and respond to antibiotics.

### Fecal microbiota transplantation shifts the gut microbiome of IBD patients

Global diversity metrics indicated robust transfer and persistent colonization of donor bacteria in patients who received a fecal transplant. From our PCoA analysis of beta diversity, we found that each patient’s samples tended to cluster (PERMANOVA *pseudo-F* = 9.55, *p* < 0.001), samples from both donors clustered together (PERMANOVA *pseudo-F* = 3.00, *p* < 0.001), and patient-patient differences drove most of the variance (Figure 1b, S1b,c). The gut microbiomes of fecal transplant recipients clearly shifted toward the communities of the donors during the trial, as indicated by the decreasing Bray-Curtis distance from donor samples, whereas those of placebo-treated patients did not (Figure 2a, S2a). This difference persisted for the ~150-day trial period, although not for every transplant patient. We categorized 3 patients receiving fecal transplants as “responders” because they showed consistent clinical, endoscopic and histologic evidence of disease improvement (Figure 2, Table S6) and categorized the other patients as “nonresponders.” Although our clinical cohort was not large enough for robust analysis between these patient populations, we did not observe a qualitative difference in donor similarity between these groups (Figure 2a).

**Fig 2.**
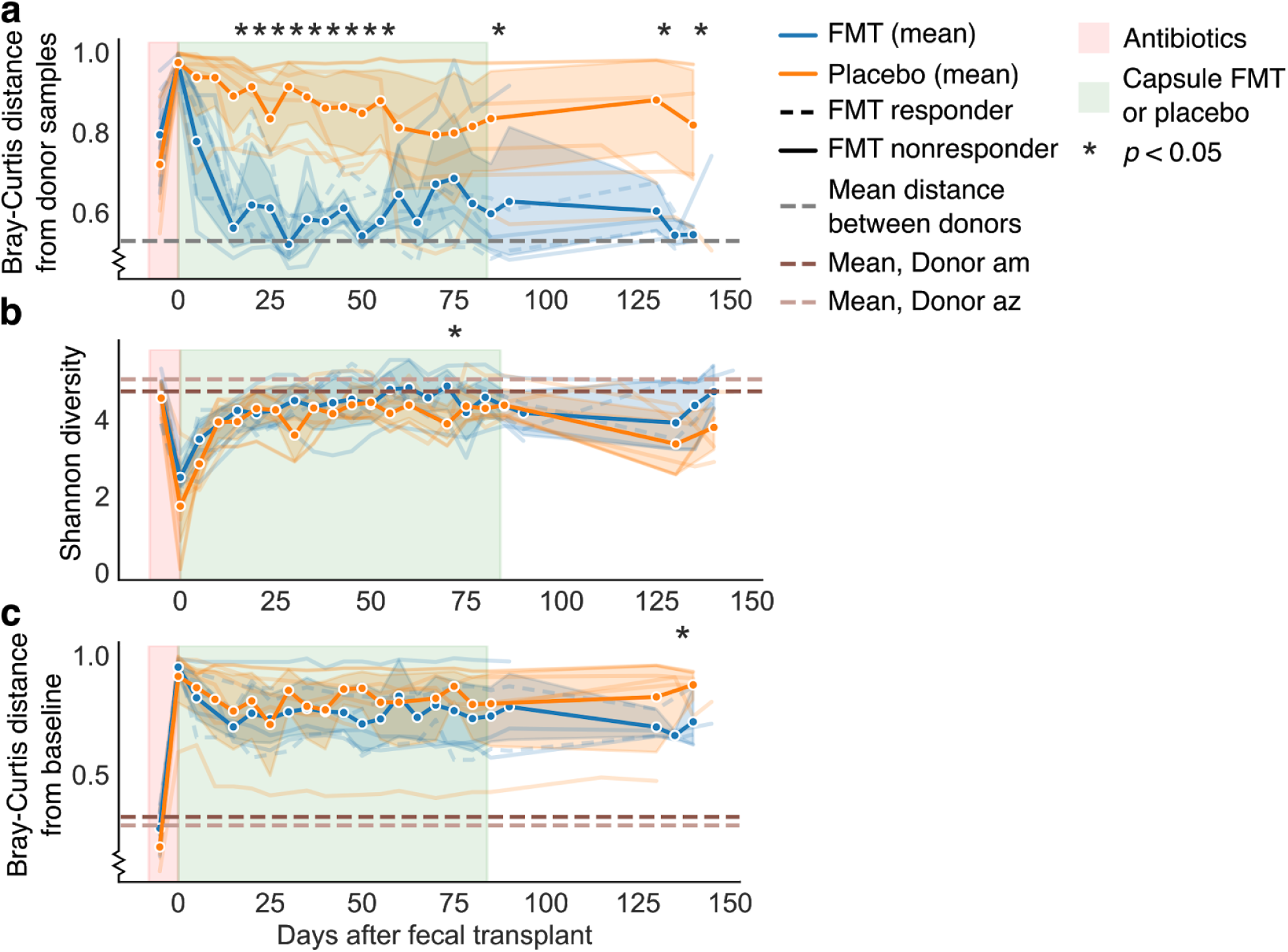
Fecal transplants resulted in global transfer and persistence of donor strains. (a) Time series of each patient’s mean Bray-Curtis distance from the donor samples. FMT patients (blue) shifted shift towards donor communities (lower values on y-axis). Bold lines and confidence intervals (95%) reflect the mean across patients. Lines for individual patients appear in the background. Lines for FMT responders are dashed, while those for nonresponders are solid. Asterisks indicate significant difference between fecal transplant and placebo patients by a Student’s *t*-test *p* < 0.05. Regions of the graph colored as in Figure 1. (b) Alpha diversity (Shannon index) of 16S profiles indicated little difference between fecal transplant and placebo recipients. (c) Similarly, the two treatment groups showed similar extents of community change when compared against their baseline samples by Bray-Curtis distance. Also see Figure S2.

Unlike previous studies, ours did not find greater alpha diversity in fecal transplant versus placebo recipients, reflecting our discovery of high microbial turnover in placebo-treated patients. Shannon index and richness in 16S and metagenomic data were similar in both treatment groups over the study period (Figure 2b, S2b–d), as was the change in bacterial community from baseline samples, according to Bray-Curtis distance (Figure 2c, S2e). These results contrast with multiple studies’ reports of increased diversity and change in gut microbiome composition in fecal transplant recipients compared with placebo recipients in diseases like *C. difficile* infection *(29, 30)*, further exemplifying how FMT can have varied effects in different diseases.

### Transferred microbial taxa exhibited varied dynamics in fecal transplant recipients

Through frequent sampling, we next profiled not only the microbial taxa that colonized transplant recipients but also their downstream dynamics, both of which likely underpin clinical response.

As in previous studies *(31, 32)*, different patients in our trial varied in their colonization rates—that is, the number and frequency of donor bacteria that successfully transferred from donor to patient (Figure 3a, Figure S3. Methods). Similar to Figure 1d, we categorized bacteria in each patient’s samples by their putative sources, including additional bacteria from the donor (“Donor”, blue lines) and shared between the donor and patient (“Shared”, grey lines) (Figure 3a). At the resolution of metagenomic bacterial species or 16S ASVs, the proportion of bacteria transferred from the donor varied between 15% and 85% of a patient’s microbiome after fecal transplant (blue lines in Figure 3a, S3a). Patients with greater numbers of donor-transferred bacteria were sometimes patients with fewer shared bacteria with the donor, but this was not always the case (e.g. FMT E and FMT P, Figure S3b). Transferred bacteria spanned phylogenetic diversity, and almost all donor bacteria from Donor am—whose stool was transplanted into four recipients—were found to colonize at least one patient (Table S4).

**Fig 3.**
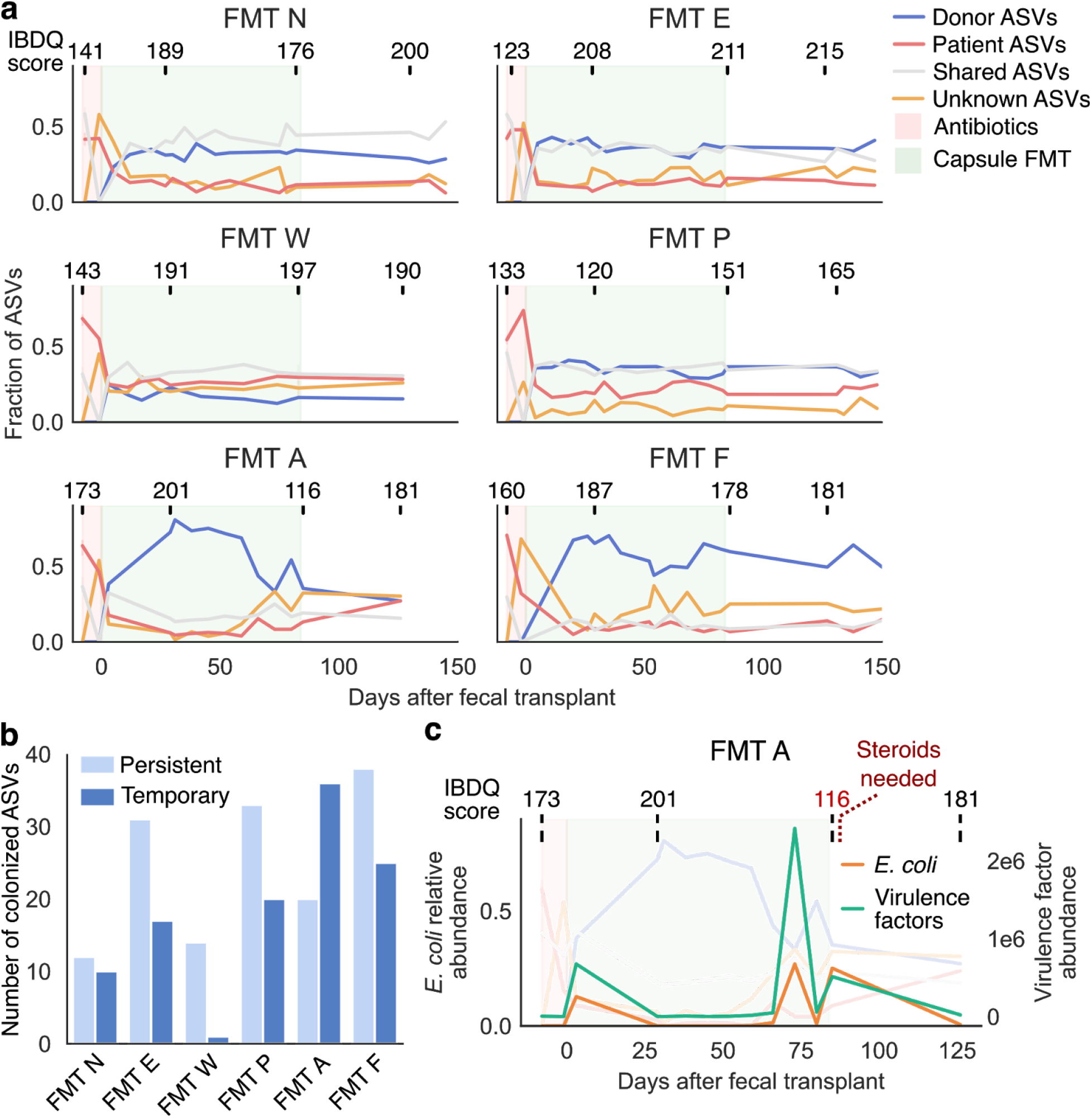
Longitudinal sampling of the microbiome revealed variable maintenance of donor bacteria. (a) Tracking bacterial sources identified bacteria transferred from donor to recipient, as well as an invasion of bacteria of unknown origin. Each time-series plot indicates the fraction of total ASVs identified as from the patient or from the donor, as shared between the two, or as from an unknown source. IBDQ scores—a standardized, clinically validated assessment tool for IBD symptoms—reported by each patient are shown above each plot. Regions of the graph colored as in Figure 1. (b) Fecal transplant recipients had varying frequencies of persistent (retained at 18-week follow up sampling after FMT) and temporary (not retained at 18-week follow up) colonization of donor bacteria. (c) After an initial period of robust colonization, patient FMT A lost a majority of transferred donor bacteria; this loss coincided with a bloom of *E. coli* and a worsening of clinical symptoms. Also see Figure S3.

Our longitudinal sampling further demonstrated that patients varied greatly in their ability to maintain colonized bacteria over time (Figure 3). Most patients showed a period of initial colonization after fecal transplantation therapy began, followed by maintenance of transferred bacteria during the daily capsule delivery period and for months thereafter (Figure 3a). In contrast, patient FMT A had robust colonization of donor bacteria early in treatment—in fact more so than any other patient—but later lost many of these bacteria (Figure 3a). These colonized-then-lost bacteria included a number of ASVs of the genera *Bacteroides*, *Faecalibacterium*, *Ruminococcus*, and others (Table S3). We categorized transferred ASVs in each patient as persistent or temporary colonizers on the basis of whether an ASV was present in follow up samples (Figure 1a). We found that different patients had different frequencies of these two types (Figure 3b, S3c). Patient FMT A clearly had mostly temporary colonizers, but even patients with largely persistent colonization had taxa that colonized only temporarily.

Patient FMT A provides an intriguing clinical case: a sharp decrease in transferred donor bacteria coincided with a bloom of *E. coli* and associated virulence factors in this patient’s gut (Figure 3c). In fact, the data suggested that the loss of donor bacteria may have preceded the bloom. Furthermore, these changes appeared to track clinical outcomes (Figure 3c). The patient reported feeling better during the early stages of treatment (week 4)—as measured by a standardized, clinically validated assessment tool (IBDQ) *(33)*—but later reported worsening of symptoms (a flare at week 12), which required administration of a steroid (prednisone) (Figure 3c). Although we can only speculate whether the *E. coli* bloom or the symptomatic change was the cause or the effect of losing donor bacteria, this case exemplifies the variability of bacterial persistence after FMT and suggests that monitoring persistence of colonizing donor bacteria may help predict FMT treatment outcomes.

### Capsule delivery from an alternate donor introduced novel taxa into an established community

By looking at the subset of patients who took capsules from an alternate donor partway through the study (Figure 1a), we were able to ask whether changing donor material resulted in additional colonization by new microbes after a new gut microbiome had been established. Although the vast majority of newly colonized bacteria came from the original donor used for colonoscopic delivery and the first month of capsules, we were nonetheless able to identify evidence of novel bacteria colonizing from the alternate donor (Figure S5). These results suggest that even after transplantation and establishment of a donor microbial community, IBD patients remain receptive to further colonization and persistence of bacteria from additional donors.

### The balance of conspecific donor, patient, and environmental strains fluctuated between dominance and parity

We then sought to profile the dynamics of individual strains within bacterial species to understand how conspecific strains (strains of the same bacterial species) from the donor, patient, and environment compete and coexist in treated patients. Previous reports have demonstrated that recipient and donor strains of the same bacterial species can coexist within fecal transplant recipients. Our longitudinal data allowed us to ask how the dynamics of this coexistence unfold over time. In particular, we asked whether donor strains could dominate over patient strains—which, because they are endogenous, may have a competitive advantage—and whether the competitive balance of strains changed over time.

We used two complementary strategies to evaluate the contributions of different strains to the resulting bacterial community in each patient: a flexible genome approach (high specificity, lower sensitivity) and a single-nucleotide polymorphism approach using StrainFinder (medium specificity, medium sensitivity) *(31)*. We focused on bacterial species with high abundance and thus sufficient sequence read depth for robust analysis (Figure 4a). Our flexible genome approach used read depth of flexible genomic regions to identify strains with identical gene content *(34)* and achieved high strain specificity by using full genome information. This approach can positively identify matches between samples with the same dominant strains (e.g., strain A versus strain A), but it cannot identify matches between samples that contain mixtures of strains (e.g., strain A versus strains A and B). Thus a “mismatch” is considered ambiguous, since the two samples might contain entirely distinct strains or a mix of shared and distinct strains, resulting in lower sensitivity. Sample comparisons therefore had three different outcomes: strain match (green in Figure 4), ambiguous (grey), and insufficient abundance or read coverage (red).

**Fig 4.**
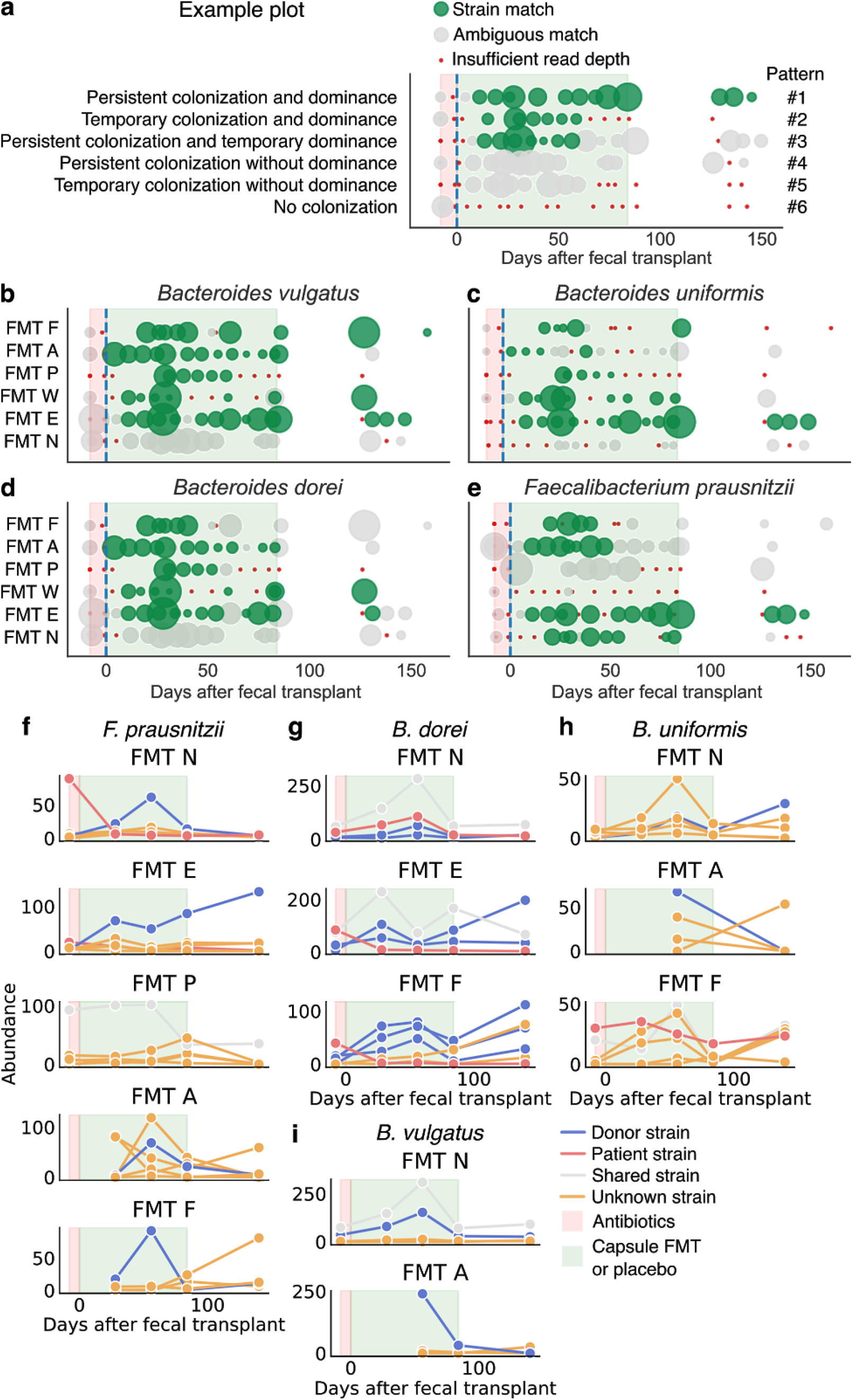
Tracking conspecific microbial strains revealed a range of competition dynamics. (a) Examples of plots for flexible genome analysis, which fell into six patterns determined by colonization and dominance (see Results). Green circles indicate a strain match between patient sample and donor sample, grey circles indicate ambiguous strain identity, and red circles indicate insufficient read depth for analysis. The size of the circle reflects the median read depth across the genome for that sample. Flexible genome plots for (b) *Bacteroides vulgatus*, (c) *B. uniformis*, (d) *B. dorei*, and (e) *F. prausnitzii*. In cases of ambiguous strain identities, we analyzed the individual contributions of strain haplotypes using StrainFinder for (f) *F. prausnitzii*, (g) *B. uniformis*, (h) *B. dorei*, and (i) *B. vulgatus.* The *y*-axes represent the frequency of different strain haplotypes for a given species, normalized by the median read depth across all marker genes for that species (see Methods). Also see Figure S4.

Using this flexible genome approach, we first confirmed that our methodology provided intuitively reasonable results by testing a number of control comparisons (e.g., donor samples did not match patient baseline samples, Figure S4). We then compared samples from fecal transplant recipients with the donor’s samples and identified many matches between donor and patient samples after the start of FMT, indicating that the dominant strain in a given patient sample was the same as in the donor (Figure 4a–d, S4c). We observed strain dynamics that fell into patterns determined by two factors: colonization (none, temporary, and persistent) and dominance (none, temporary, and persistent). These factors resulted in six observed patterns, and we provide examples of these patterns in Figure 4a:

1. Persistent colonization and dominance: The donor strain colonized after a fecal transplant, came to dominate the strain community, and persisted as the dominant strain throughout the trial.
2. Temporary colonization and dominance: The same as #1, but all strains were subsequently lost.
3. Persistent colonization and temporary dominance: The same as #1, but at later time points, another strain from the patient or from an unknown source became equally or more abundant than the donor strain.
4. Persistent colonization without dominance: The donor strain colonized and persisted in the patient but never dominated the patient’s microbial community.
5. Temporary colonization without dominance.
6. No colonization.

We observed examples of each of these patterns, indicating the range of competitive dynamics between strains that can unfold after fecal transplant. Many *Bacteroides* species were successful in persistently and dominantly colonizing (pattern #1), including *B. vulgatus* (patients E, W, F), *B. dorei* (patients E, W), and *B. uniformis* (patient E) (Figure 4b–d). The same bacteria in other patients persistently colonized but only temporarily dominated the patient community (pattern #2), including *B. vulgatus* in patient P, *B. dorei* in patients P and F, and *B. uniformis* in patients W and P. One patient (N) appeared to be more resistant to colonization by donor *Bacteroides* species, nearly always showing a mix of strains (patterns #4 or #6). We also observed temporary colonization and dominance for *B. ovatus* and *B. caccae* in fewer patients (Figure S4c). With regard to *F. prausnitzii*—a commensal bacterium thought to be related to gut health and negatively correlated with IBD *(24)*—some patients exhibited persistent colonization and dominance of a donor strain (pattern #1, patient E), others were only temporarily dominated by the donor strains (pattern #3, patients P, N), and still others adopted strains that came from both the donor and unknown sources (Figure 4e).

In cases of ambiguous matches, our flexible genome analysis could not define the contributions of individual strains from different sources, leaving some competitive dynamics undefined. For example, a donor strain of *B. dorei* temporarily dominated the community of Patient F, but later strain matching gave ambiguous results, which could mean that the donor strain disappeared from the community or that it coexisted with another strain (Figure 4d). Consequently, to observe individual contributions of strains in mixed communities, our second approach reconstructed SNP haplotypes using StrainFinder, allowing us to distinguish contributions of strains from donor, recipient, and unknown sources over time. Although the sensitivity of StrainFinder allows us to quantify individual strains, its dependence on marker genes makes it less specific than our flexible genome approach *(31)*. Because StrainFinder requires high sequencing depth to properly model strains, we combined longitudinal samples (*n* = 1–5 samples) into five time points (Methods, Figure 4).

We found that donor and patient strains frequently coexisted, even after temporary dominance by the donor strain. We first confirmed that the results from StrainFinder aligned with our flexible genome analysis, as shown for *F. prausnitzii* in FMT N and FMT E (Figure 4e). In the case of FMT A and FMT F, we observed that by the end of the clinical trial, strains of unknown origin largely displaced those from the donor (Figure 4e). The dominant strains of some species (*F. prausnitzii* in FMT P, *B. dorei* and *B. vulgatus* in FMT N) were shared between the donor and patient (Figure 4f–i). If we assume that it is unlikely that two unrelated individuals carry identical strains, these results indicate that even high-resolution methods like StrainFinder depend on marker genes and cannot always resolve unique strains. For *B. dorei* in FMT F and *B. uniformis* and *B. vulgatus* in FMT A, donor strains temporarily dominated, with strains of unknown origin appearing in the patients’ follow-up samples (Figure 4g–i). We observed a similar variety of competitive dynamics in other abundant species (Figure S4f–i).

Taken together, these results demonstrate not only that donor and recipient strains can coexist *(32)*, but also that the balance of this coexistence changes over time. In many cases, donor strains were able to outcompete endogenous patient strains, but this dominance was dynamic, with donor and patient strains often competing with strains from unknown and possibly environmental sources later in the trial.

### Fecal transplants transferred beneficial microbial functions that varied across time

Beyond the microbes themselves, the functions those microbes perform in the gut ecosystem may be important to restoring gut health. Therefore, we tracked colonization by functional genes implicated in maintaining health. We observed the transfer from donor to patient of genes involved in complex carbohydrate metabolism (Figure 5a, S6a), mucin digestion (Figure 5b, S6b), and butyrate production (Figure 5c, S6c), and many of these genes persisted in their new hosts. These transferred genes included genes from health-associated commensals like *F. prausnitzii* and *Bacteroides* (Table S5) *(24, 35)*. Like the microbes themselves, these functional genes were transferred at different rates and had variable dynamics across patients (Figure 5), and these dynamics largely mirrored patterns of microbial colonization (Figure 3).

**Fig 5.**
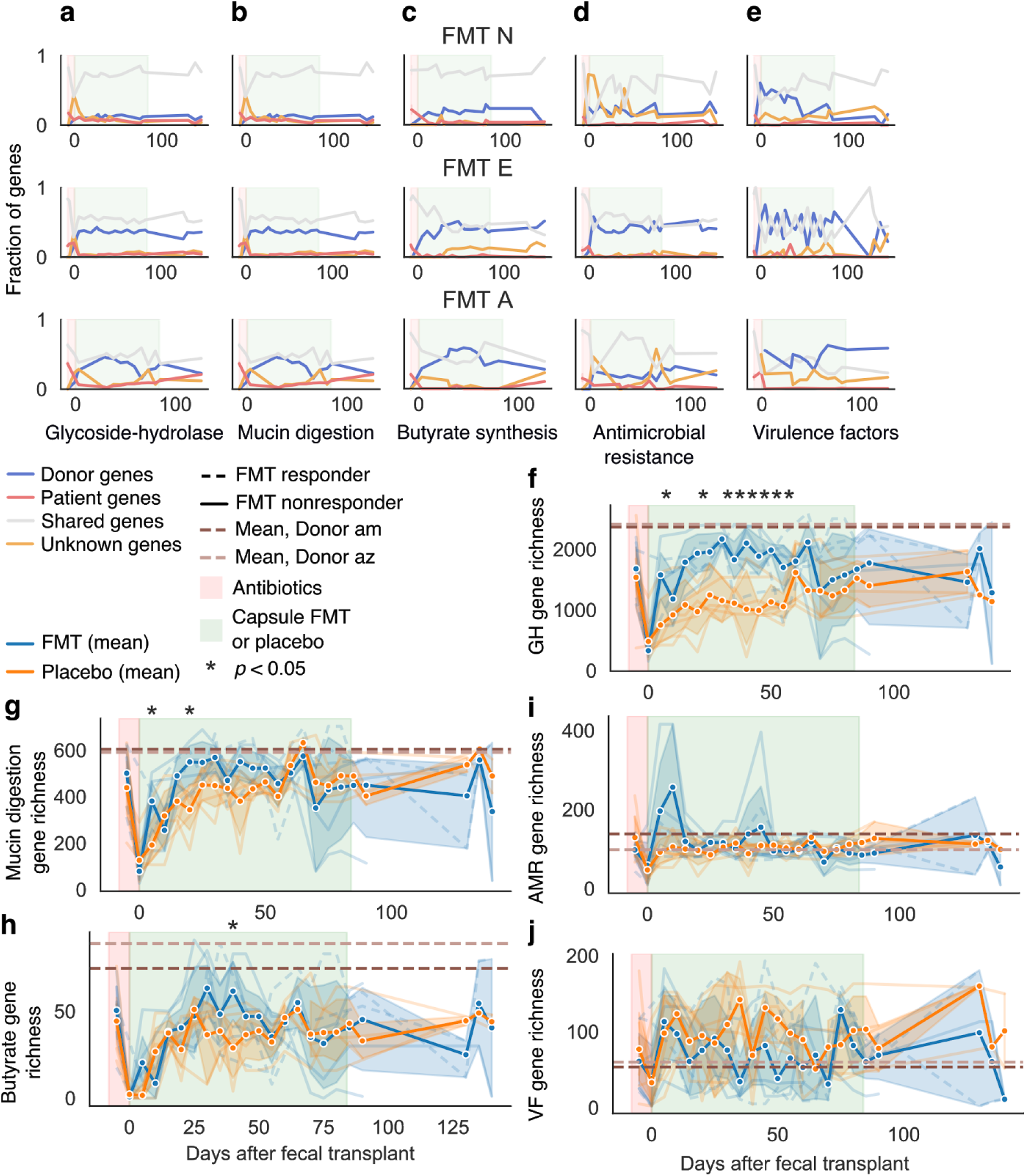
Bacterial functions also transferred across human hosts. We tracked the transfer of bacterial functions using shortBRED and curated databases (see Methods). We observed colonization and persistence of functional genes in transplant recipients for (a) glycoside hydrolase (GH), (b) mucin degradation, (c) butyrate biosynthesis, (d) antimicrobial resistance, and (e) virulence factor genes. Shown are plots for three of our transplant recipients (see Figure S6). We also examined overall diversity of these genes compared with placebo-treated patients. Fecal transplant recipients showed higher diversity of (a) glycoside hydrolase genes, but not of (b) mucin degradation genes, (c) butyrate biosynthesis genes, or (d) antimicrobial resistance genes. (e) Placebo patients had slightly higher diversity of virulence factors, but the difference was not significant. Asterisks indicate significant difference between fecal transplant and placebo recipients by a Student’s *t*-test *p* < 0.05. Also see Figure S6.

Overall, fecal transplant recipients showed greater richness in genes involved in complex carbohydrate metabolism than did placebo-treated patients, suggesting increased capacity to digest dietary polysaccharides (Figure 5f). On the other hand, fecal transplant recipients showed similar levels of butyrate biosynthesis (figure 5g) and mucin digestion genes (Figure 5h). So although donors were chosen for high butyrate production—reflected in high diversity of butyrate biosynthesis genes in donor samples (Figure 5c, dashed lines)—fecal transplant did not result in wholesale transfer of these genes; transplant recipients had similar butyrate gene diversity compared with their baseline samples and with placebo-treated patients. Two fecal transplant recipients—but no placebo-treated patients—reached comparably high butyrate gene diversity, but these changes were temporary. Thus, fecal transplants effectively transfer beneficial microbial functions across hosts but may not result in persistent colonization or an overall increase in the diversity of genes related to those functions.

### Fecal transplants also transferred antibiotic resistance and virulence factors

Not all microbial functions transferring to fecal transplant recipients are beneficial. Although, clinically, fecal transplants can effectively clear patients of antibiotic-resistant infections, there is an ongoing debate on whether the therapy could introduce novel resistance genes, with negative clinical effects *(15)*. We observed the transfer from donors to patients of numerous antimicrobial resistance genes—including those for resistance to all major classes of antibiotics—and many of these genes were maintained for the full trial period (Figure 5d, S6d, blue lines). Nevertheless, we found that the resulting diversity and abundance of antibiotic resistance genes in fecal transplant recipients was not heavier than in our healthy donors (Figure 5i, S1g,h) and that transferred resistance genes were generally outnumbered by endogenous ones (Figure 5d, S6d). Thus, although the transfer of resistance genes is an unavoidable result of the complexity of this therapy, no clinical or bioinformatic evidence indicates that fecal transplants increase the overall risk of antibiotic resistance. Indeed, there are many case reports suggesting that FMT can help rid patients of antibiotic resistant infections *(36, 37)*.

Similarly, we observed the transfer of virulence factors. Despite a lower incidence of virulence factor genes in donors than in patients (Figure 5j), we observed colonization and persistence of such genes, which made up a significant portion of the virulence factor pool (Figure 5e, S6e, blue lines). Many patients exhibited an increase in the abundance of virulence factors during or shortly after antibiotics (Figure S6f). Two patients had an inordinate burden of virulence factors, including the patient FMT A who had a bloom of *E. coli*, and patient Placebo V (Figure S6g). Patient Placebo V’s health declined during the trial (Table S5), and this patient had one of the most dramatic microbial turnovers in response to antibiotics (Figure 2d). This turnover resulted in a microbiome dominated by newly acquired Proteobacteria and associated virulence factors from unknown sources, again highlighting the risks of antibiotics in IBD patients (Figure S6h). In sum, although we observed the transfer and persistence of donor-derived virulence factors and antibiotic resistance genes, these transfers were modest and often outweighed by the endogenous microbial community’s own virulence and resistance. *IgA coating of gut microbes identified shared immune responses to commensal and IBD-associated bacteria*

In the context of IBD, interaction with the host immune system is perhaps the most important microbial function. Although it is reasonable to expect that transferring microbes would also transfer their endogenous metabolic capacities, it is much less certain whether transferred bacteria will elicit similar immune responses—particularly adaptive immune responses—in a new host with a different immune system.

To understand host response to transferred gut bacteria, we used immunoglobulin A sequencing (IgA-seq) to profile bacteria coated with IgA antibodies *(17)*. Secretory IgA is the primary antibody of mucosal surfaces, including the gastrointestinal, respiratory, and urinary tracts *(38)*. Thought to act primarily by blocking proteins on the surface of invading pathogens, IgA has more recently been suggested to play a role in facilitating mucosal colonization by commensal bacteria *(39, 40)*. Although bacteria interact with the host immune system in many ways (e.g., via excreted metabolites *(12)* or direct contact with the epithelium *(16)*), we used IgA-seq as a proxy for bacterial immune function/interactions because it is one of the few immune functions that can be measured *in vivo* and at high-throughput for bacteria in stool samples. To identify IgA-coated and -uncoated bacteria, we used fluorescence-activated cell sorting (FACS) to separate these fractions of gut microbiome samples, and we 16S-sequenced the fractions (~50,000 cells per fraction) to a median depth of 165,000 reads. We calculated IgA coating scores for each bacterium as the log-fold change in abundance of that bacterium in the IgA-coated and IgA-uncoated fractions, such that a positive value indicated high IgA coating and a negative value indicated low IgA coating. To quantify similarity in IgA coating between samples, we calculated the Pearson correlation of IgA coating scores across all bacterial ASVs.

If IgA coating of different bacteria were the same across all patients and donors, then the likelihood of successful transfer of IgA coating would be very high. We therefore first asked whether overall IgA coating of bacterial ASVs differed among patients. We found that IgA coating scores of bacteria in two samples taken at different time points from a single healthy donor had high agreement (Pearson *r* = 0.7, *p* < 1e^−25^) (Figure 6a), suggesting that IgA coating of bacteria in healthy individuals is stable across time; this result gave us confidence in our methodology. We then compared the mean IgA coating in Donor am’s samples versus the mean IgA coating scores across all patient’s baseline samples and found moderate correlation (Pearson *r* = 0.43, *p* < 1e^−9^, Figure 6b). The strength of this correlation (donor samples versus baseline samples) for individual patients, however, varied considerably (Pearson *r* of 0.1–0.7, Figure S7a,b). This result indicated that bacterial IgA coating can vary greatly across patients and raised the prospect that transferred bacteria may not retain their immune function. Some studies have suggested that IgA might target blooming or abundant bacteria to promote homeostasis *(41, 42)*, but we did not observe a correlation between a bacterium’s IgA coating and its abundance, variance, or bimodality (Fig S7c–e).

**Fig 6.**
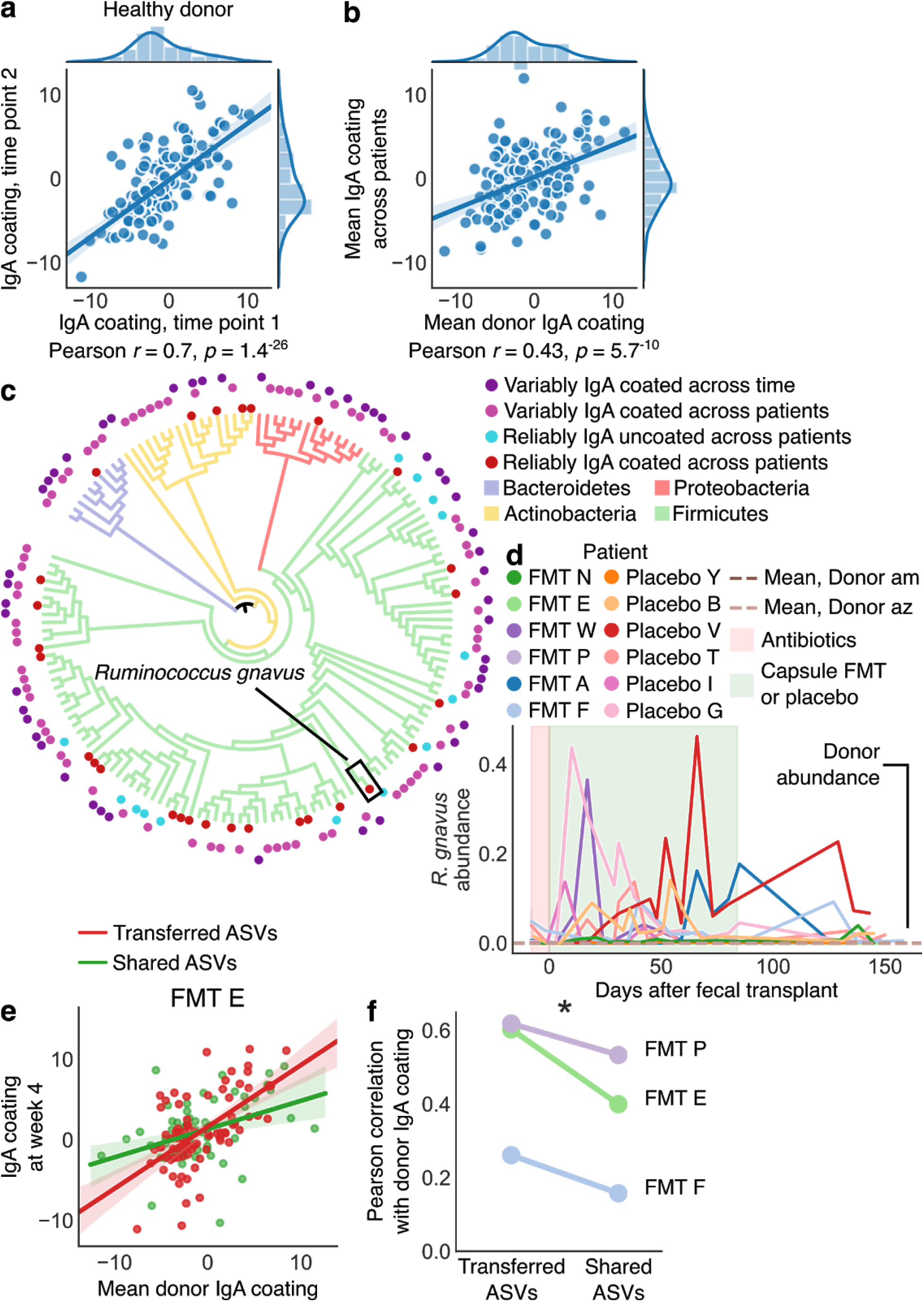
IgA-coating of the gut microbiome revealed broad patterns of microbiome IgA coating and host-specific IgA responses. IgA enrichment of different bacteria was (a) highly correlated across different samples from a healthy donor and (b) even well correlated between the same donor and the patients’ baseline samples. In panels (a) and (b), each point is a bacterial ASV, and the x and y axes represent the IgA coating score of that ASV in different samples. (c) A phylogenetic tree containing bacteria that were identified as reliably IgA coated across patients (inner red circles), IgA uncoated across patients (blue circles), or variably IgA coated/uncoated either across patients or within a patient across time (outer purple circles). Tree branches are colored by phylum. (d) Relative abundance of *Ruminococcus gnavus* in our patient cohort. (e) Correlations of IgA enrichment in shared and transferred bacteria indicated stronger correlation in transferred bacteria, which are more likely to be exact matching strains. (f) This pattern was observed in all patients receiving transplants from Donor am. Also see figure S7.

We then sought to identify bacteria that were strongly IgA coated or uncoated across all patients and samples, because these bacteria likely trigger similar immune responses when transferred across hosts. Indeed, we found 28 bacterial ASVs (*p* < 0.0001 by a permutation test) that were reliably and strongly IgA coated across patients and within patients through time, including before and after FMT (see Methods). These bacteria were a phylogenetically diverse group comprising organisms from all major phyla (Figure 6c, Table S6), including known commensals (e.g., *Bacteroides*) and many taxa that transferred from donors to fecal transplant recipients (e.g., *Bacteroides*, *Lachnospiraceae*, Table S6). We also identified some Proteobacteria (including known opportunistic pathogens) and *Ruminicoccus gnavus*—as well as the closely related *R. torques—*as reliably IgA coated in patients and donors (Figure 6c, S8, Table S6). Although we did not observe *R. gnavus* transfer from donors to patients, we found significant blooms of this bacterium (up to 40% relative abundance) in our patient cohort, while it was essentially absent in our donors (Figure 6d). This finding is in keeping with those of other research groups, who have reported that *R. gnavus* blooms specifically in patients with IBD *(43)*. These results further suggest that *R. gnavus* may play a role in immune dysregulation in IBD and warrants additional study.

Bacteria that were reliably uncoated by IgA (*n* = 14, *p* < 0.0001 by a permutation test) across patients were less phylogenetically diverse and tended to represent known commensals. All of these bacteria were Firmicutes, including known butyrate producers like *Faecalibacterium*, *Alistipes*, *Roseburia*, *Oscillibacter*, *Butyricicoccus*, and other *Lachnospiraceae* species (Figure 6c, Table S6). On the basis of these results, we hypothesize that being uncoated by IgA may be more specific than being coated. Together, these data suggest that bacteria eliciting strong or negligible IgA responses across individuals likely play important roles in regulating host gut immunity and that transferring these bacteria can replicate microbial immune function in a new host.

### Strain specificity revealed the transfer of host immune function in fecal transplant

We observed a final category of diverse bacteria that were sometimes IgA coated and sometimes uncoated across different individuals or even within the same individual across time (Figure 6c, S8, Table S6). If confirmed, such variable immune effects across patients could complicate FMT treatment, which currently assumes the transfer of microbial functions from donors to patients. If the same microbe can trigger an IgA response in one person while suppressing it in another—which may lead to proinflammatory or anti-inflammatory responses *(44)*—then FMT may be less effective and less predictable than assumed at transferring immune function.

We hypothesized that variable IgA responses could stem from two processes: (1) divergent host immune responses or (2) strain specificity of IgA coating. If the second process were responsible, a given bacterium would be IgA coated or uncoated because each patient would be responding differently to different strains of bacteria with identical 16S sequences, the genetic marker for our IgA-seq data. But each of these patients would nevertheless respond in the same way to the exact same strain, and thus transfer of the same strain across patients would transfer a similar IgA response.

To establish the role of strain specificity in explaining variably IgA-coated bacteria, we examined two bacterial subsets in fecal transplant recipients: bacteria that were shared by donor and patient and bacteria that transferred from the donor to the patient after a fecal transplant (Methods). Thus, for each bacterial 16S sequence, the first subset included a potential mix of donor and patient strains, while the second subset likely contained only one strain from the donor that then transferred to the patient. We found that the subset of strains transferred from the donor (exact strain matches) had higher correlations of IgA coating scores than bacteria shared between the donor and recipient (mixed strains) (Figure 6e); this pattern held across three patients who received fecal transplants from a single donor (Figure 6f).

These results suggested that exactly matching strains triggered more-similar immune responses than mixed strains with identical 16S sequences, demonstrating that variable IgA coating of bacteria could in part be explained by strain specificity. Although in one patient, the correlation of IgA coating scores of exactly matching strains was still weak (Pearson *r* = 0.3), this finding further established that immune functions of transferred microbes can be broadly replicated in fecal transplant recipients, bolstering the prospects of engineering the gut microbiome to modulate host immunity and disease.

## Discussion

### Disease context shapes gut microbiome recovery after antibiotics

We found that administering broad-spectrum antibiotics destabilized the gut microbiome of placebo-treated IBD patients. Unlike healthy patients in other studies, our IBD patients readily lost and did not recover their original gut bacteria, exposing these patients to colonization by new bacteria, which in some cases ultimately outnumbered endogenous species. Previous work has reported that the microbiomes of IBD patients are more variable than those of healthy people *(5)*, and our results establish that interventions like antibiotics can exacerbate this instability. Given that multiple clinical trials have studied the use of antibiotics to treat IBD *(45–50)*, our results raise a red flag about potential unintended consequences of antibiotics for patients with IBD or other microbiome-related diseases. Further assessment of the effects of different antibiotics on the gut microbiome of IBD patients would improve clinicians’ ability to make informed clinical decisions about antibiotics’ risks.

### Microbial strains exhibit a range of colonization dynamics

Our results illustrated a range of microbes and functions that can persistently colonize fecal transplant recipients. Only a small subset of rare bacteria appeared to never transfer (Table S5), suggesting that essentially all bacteria can be transferred between people. This finding reaffirms that the gut microbiome can be clinically engineered by transplanting whole gut microbial communities. It remains unknown whether more-targeted therapeutics using synthetic communities will show the same ability to colonize recipient hosts.

For fecal transplants or other targeted microbial therapeutics to have a clinical effect, colonized microbes must persist at sufficient abundance. We observed a variety of fates of colonizing microbes in each patient over time—from wholesale colonization and persistence of new taxa to fleeting passage of individual strains. A significant fraction of microbes that colonized the recipient for multiple weeks later disappeared, suggesting that even though many bacteria can colonize a patient temporarily, competition, nutritional requirements, or immune system interactions may hamper persistence. This problem might be addressed by administering fecal transplants along with other treatments aimed at maintaining colonized bacteria (e.g., prebiotics, maintenance therapy) *(42, 51, 52)*.

Coexisting conspecific strains also showed a range of competitive dynamics: in some patients donor strains dominated endogenous strains while in others, endogenous strains remained more abundant. Competitive dynamics like these may contribute to variable clinical responses to whole-gut-microbiota transplantation and are likely to play an even greater role in more-targeted microbial therapeutics, whose efficacy hinges on the dynamics of a small number of strains.

In addition, Patient FMT A offered an example of how patient health can affect the colonization of donor bacteria. After more than a month of stable colonization, this patient lost a large portion of transferred strains in a short period, which coincided with—or potentially preceded—a bloom in pathogenic bacteria and severe worsening of symptoms. This dramatic decline warns us that continued patient monitoring may be needed to maintain treatment efficacy, particularly with chronic diseases like IBD. Patients who lose colonized donor bacteria could be retreated, restarting the clock on donor-strain persistence and intended clinical effect. Furthermore, although it remains unknown which of these shifts came first—loss of donor bacteria, bloom of pathogens, or worsening of symptoms—the progressive unfolding of these events raises the possibility that real-time tracking of patient microbiomes may enable early intervention and prevention of IBD flares.

### Microbial and immune functions transfer across human hosts

We found that specific beneficial functions transferred from donors to patients and could also persist. Many gut microbiome studies have focused on the benefits of butyrate production, for example, and we were able to track the transfer of butyrate production genes from donor to patient. But even after receiving new butyrate genes from a donor, fecal transplant recipients did not show higher butyrate gene diversity compared with placebo-treated patients. This observation suggests that it may be difficult to increase overall genetic capacity for butyrate production via fecal transplants. Of course, the diversity of genes related to a function does not necessarily reflect the activity of those biochemical pathways. Instead, it may be more fruitful to focus on which butyrate-producing organisms are present (are some microbes more productive than others?) and which nutrients (e.g., dietary fibers) are available to those bacteria. Examining how colonization, persistence, and environmental context alters the activity of transferred gut bacteria will bring us closer to understanding the pharmacokinetics of gut microbiome engineering.

We found that fecal transplants can also transfer unintended functions (e.g., antibiotic resistance genes, virulence factors). To date, no evidence suggests that such unintended transfers have appreciable clinical effects *(1, 53)*, but possibilities must be considered, particularly since antibiotic resistance can transfer between gut bacterial species *(53, 54)*. Although it is probably impossible to purge an intact fecal community of all antibiotic resistance, targeted microbial therapeutics may be able to minimize or avoid it.

In the context of IBD, the function most critical to transfer and persist in the patient is the gut microbiome’s immune function. Our identification of numerous reliably and strongly IgA-coated or -uncoated bacteria across all patients and donors indicated retention of immune function across hosts. Strongly IgA-coated bacteria included IBD-associated bacteria (*R. gnavus*, *E. coli*), as well as known commensals (*Bacteroides*, *Blautia*)—a finding that complicates the frameworks of research suggesting that IgA-coated bacteria are largely pathogenic and inflammatory *(17, 38)*. It may be that IgA coats any bacterium colonizing the gut mucosa, whether friend or foe. This speculation fits with recent reports of *Bacteroides* commensals using IgA to colonize the mucosa *(39)* and with the observation that IgA and gut bacteria tend to concentrate in the outer layers of mucus in mice *(55)*.

Furthermore, we establish that IgA coating of bacteria can be transferred across human hosts, suggesting that transferring gut microbes may be broadly effective in triggering specific and nonspecific IgA coating and immune pathways. In addition to bacteria that were reliably IgA-coated or -uncoated, many bacteria were variably IgA coated across patients, suggesting that potential host specificity of immune function could complicate clinical responses to fecal transplants. We found that this variability was in part due to the strain specificity of IgA coating: strains that transferred from donor to patient tended to have similar patterns of IgA coating. This specificity seems to contrast with previous reports of polyreactive IgA activity in the mouse small intestine *(56)*. It is highly unlikely that this signal resulted from the IgA coating of bacteria in the daily capsules because of their small volume (one capsule per day) and the necessity for those bacteria to pass through the small and large intestine.

It is further possible that a donor’s immune context may play a role in the transfer of IgA coating from donor to patient, either because of donor-specific immune responses or other donor-specific factors like diet or microbial community. For example, a particular bacterial strain might express different surface receptors depending on the nutrients in a host’s diet, which may then alter what would otherwise be identical immune interactions *(44)*. We further speculate that an IgA-coated bacterium from a donor, transferred into a patient, may retain its IgA coating, not because the patient innately coats that bacteria, but because the patient’s immune system learns the coating pattern from the donor’s IgA. In such a scenario, IgA coating and immune function may display an “inertia” when transferred between hosts. Additional study into the immune factors that generate transferable and variable IgA responses will illuminate our ability to manipulate host immunity via the gut microbiome.

In summary, our study offers a first look at the dynamics of colonization and persistence of microbes, their metabolic functions, and their immune functions in IBD patients treated with FMT. Our dense time series sampling revealed surprising complexity in microbial transfer and emphasized that for chronic diseases like IBD, continuing patient care may be necessary to maintain newly colonized bacteria. Our observations of broad transfer of microbes and their functions further demonstrate the power of FMT to alter a patient’s gut microbiome and set the stage for developing targeted drugs that introduce and maintain specific microbes and functions to treat disease.

## Materials and Methods

### Clinical cohort and sample collection

We collected samples from a clinical cohort recruited at the University of Vermont Medical center in Burlington, Vermont, USA. Patients collected semiweekly stool samples at home or in the clinic, storing samples in RNAlater solution (ThermoFisher) and mailing them to a processing facility at OpenBiome in Somerville, MA. During clinical evaluations at University of Vermont Medical Center, we also collected fresh stool samples at baseline and at 4, 12, and 18 weeks after the initiation of FMT; we stored these samples in a glycerol buffer (1X PBX, 25% glycerol, 0.05% L-Cysteine).

### DNA extraction and sequencing

We triple-washed RNAlater from samples in 1X PBS and extracted DNA using a MoBio Powersoil DNA extraction kit. 16s rDNA libraries were prepared and sequenced by the Broad Institute Genomic Platform, using the Earth Microbiome Project protocols and paired-end 250-base-pair reads on an Illumina MiSeq *(57)*. Shotgun metagenomic libraries were likewise prepared by the Broad Institute using Nextera protocols and sequenced on an Illumina NextSeq.

### IgA sequencing

Samples were processed as described previously *(17)*. We centrifuged glycerol-stored stool samples at 50 × g at 4°C for 15 min and then washed them three times in 1 mL PBS/1% BSA at 8,000 × g for 5 minutes. We collected the presort fraction as 20 μL after resuspension before the final wash and stored the washed samples at −80°C. We then resuspended the cell pellet in 25 μL of 20% Normal Rat Serum (Jackson ImmunoResearch) in PBS/1% BSA and incubated the samples for 20 min on ice. After incubation, we added 25 μL 1:12.5 α-mouse-IgA-PE (eBioscience; clone mA-6E1) to each sample and incubated samples on ice for 30 minutes. Finally, we washed samples three times in 1 mL PBS/1% BSA, resuspended them in PBS/1% BSA, and transferred them to blue filter cap tubes (VWR 21008-948) for flow sorting. We sorted an average of 50,000 cells from the IgA-positive and IgA-negative bacteria into sterile microcentrifuge tubes on the BD FACSAria II at the MIT Koch Institute Flow Cytometry Core (Cambridge, MA). We then centrifuged the samples, removed the supernatants, and resuspended the pellets in a final volume of 10 uL sheath fluid. Samples were stored at −80°C until DNA library prep, in which 2 uL (~10,000 cells) was used directly as the template for PCR.

### Data analysis

We analyzed 16S data using Qiime2 *(25)*, DADA2 *(26)*, and custom Python scripts. We assigned taxonomic labels to 16S sequences using the SILVA database *(58)*. We quantified the abundance of microbial species from shotgun metagenomic sequencing using MetaPhlAn2 *(27)*. To visualize changes in alpha and beta diversity, we calculated the mean values of samples within five-day windows, and compared these values across treatments using a Student’s t-test.

To track the sources of various bacteria, we defined all bacteria observed in any of a patient’s baseline samples and in the donor sample as “Shared,” all other bacteria present in baseline samples as “Patient,” all bacteria absent from baseline but shared with the donor as “Donor,” and finally all others as “Unknown.” We defined “persistent” colonization as a bacterium (ASV or metagenomic species) that transferred exclusively from a donor and appeared in at least three samples after the initiation of FMT and remained present in at least one follow-up sample at ~18 weeks after initial transplant. We defined “temporary” colonization similarly, except that such bacteria did not appear in any follow up samples.

Assigning sources using this strategy has its limitations. Because many strains within common bacterial taxa have identical 16S sequences, 16S-based techniques may register many unique strains are identified as a single ASV. For example, in the case of *E. coli*, all transplant and placebo-treated patients and the donors shared a single *E. coli* ASV, but it is highly unlikely that every patient in fact shared the same *E. coli* strain. Thus our strategy may occasionally falsely identified the source of a given ASV. Consequently, we focused on the overall frequency (occurrences) of ASVs from different sources, instead of the abundance of each ASV. This approach minimizes the signal from highly abundant but potentially incorrectly identified ASVs, as it weights all ASVs equally.

To quantify the transfer of bacterial functions, we used ShortBRED *(59)* to determine the abundances of genetic functions of interest, including butyrate biosynthesis *(60)*, mucin degradation *(61)*, glycoside hydrolase activity *(62)*, antimicrobial resistance *(63)*, and virulence factors *(64)*. We quantified the abundance of quinolone resistance in baseline samples and in the 10 days immediately after antibiotics stopped, and we compared these abundances using a Student’s paired *t*-test. We also calculated the area under the curve and compared these values using a Student’s paired *t*-test. We visualized the abundance of genetic functions in FMT and placebo patients using the same five-day windows as described above. We identified the sources of antibiotic resistance genes and virulence factors in the same way we identified as bacterial sources.

To quantify the transfer of strains, we used two strategies: one based on flexible genome content and the other on single-nucleotide variants (SNVs). For the first, we used a strategy similar to that described previously *(34)*. Briefly, we mapped metagenomic reads from each sample to reference genomes for each species using BWA *(65)* and quantified the number of reads mapping to each unique 1000-bp segment of the reference sequence. To compare the strains in two samples, we then compared the read depths in each sample across all 1000-bp segments. We identified a strain match as those comparisons for which no segment with a read depth greater than the median for that sample was entirely absent from the other sample (Figure S4). Comparisons that did not meet these criteria were called ambiguous. Comparisons where either sample had a median read depth less than 5 were not considered because of insufficient abundance and read depth. To reconstruct the individual contributions of strain haplotypes, we used StrainFinder *(31)*. To build the input alignments for StrainFinder, we used BWA *(65)* to align metagenomic reads from each sample to a database of AMPHORA genes *(66)*—a set of single-copy, universally carried bacterial genes—from various gut bacteria. We used SAMtools *(67)* to tally the nucleotide identities found at each position and filtered stringently to remove reads with poor mapping quality, rare alleles, and sites with inordinate read depth. To provide greater depth of reads to StrainFinder’s maximum-likelihood model, we combined reads from samples across our time series as described above. We considered only genomes with a median read depth exceeding 50 in at least two samples. StrainFinder outputs the relative contributions of different strains to the abundance of a given species, so we normalized these values using the median read depth in each sample to better reflect the relative abundances of each strain across samples.

To understand host-immune interactions, we analyzed 16S data from IgA-seq using Deblur *(68)*. We calculated IgA coating scores as the log2-fold change between the IgA-coated and -uncoated fractions. We observed that IgA coating scores somewhat followed a normal distribution (Figure S7f); thus, for each sample, we categorized strongly IgA-coated or -uncoated bacteria as those bacteria that were greater than or less than the mean +/– 1 standard deviation. To identify bacteria that were reliably IgA coated or uncoated across all samples or across all patients, we used a one sample *t*-test, with an FDR-adjusted *p* value of < 0.1. We further evaluated whether or not these reliably IgA-coated or -uncoated ASVs were statistically significant by a permutation test, in which we randomly shuffled the ASV labels in each sample 10,000 times and counted the occurrences of the same or greater number of reliably IgA-coated or -uncoated ASVs. We constructed a phylogenetic tree of 16S sequences using FastTree *(69)* and visualized it using iTOL *(70)*.

## Acknowledgments

We thank the people at OpenBiome for their collaboration on this clinical trial. We are grateful to the BioMicroCenter at MIT and Microbial Omics Core at the Broad Institute for their assistance with library preparation and sequencing. We thank Ellen W. Chu for substantive and editorial feedback on the manuscript.

## Funding

This work was supported by funding from the Center for Microbiome Informatics and Therapeutics. We also thank the National Science Foundation GRFP for supporting NDC. Funders had no role in study design, data collection and analysis, decision to publish, or preparation of the manuscript.

## Author contributions

NDC, JWC, LTTN, MBS, ZK, PLM, EJA designed the study; JWC and PLM coordinated patient recruitment, treatment, and sampling; MBS and ZK coordinated fecal microbiota transplant production and delivery to the clinic; NDC, JWC, LTTN, MBS, ZK coordinated sample collection; NDC, LTTN coordinated data generation; SMK performed IgA-seq experiments; NDC analyzed the data; and NDC wrote the first draft of the manuscript with substantive input from all authors.

## Data and materials availability

We deposited raw sequence files of 16S, metagenomic, and IgA-sequencing results in NCBI’s SRA database (BioProject PRJNA475599). Metadata is included as supplementary tables of this manuscript. Additional 16S ASV tables and metagenomic species tables as well as code used to generate the figures and analyses can be found at https://github.com/nathanieldchu/uc_fmt.

## Competing interests

EJA is a cofounder and shareholder of Finch Therapeutics, a company that specializes in microbiome-targeted therapeutics. PLM is a Medical Director at Takeda Pharmaceutical Company. MBS is a cofounder and CEO of Finch Therapeutics. ZK is a cofounder and Executive Vice President of Finch Therapeutics.

## Supplementary Materials

**Fig S1.** Related to Figure 1: instability in the gut microbiome of IBD patients after antibiotics. (a) Turnover in gut microbiome species in healthy patients administered a course of the broad-spectrum antibiotics vancomycin, gentamicin and meropenem *(28)*. (b) PCoA of unweighted UniFrac distance of 16S. (c) PCoA of Bray-Curtis distances based on metagenomic species. (d) PCoA trajectories, based on metagenomics species, of placebo patients. (e) Source plots of metagenomic species in placebo patients. (f) Abundance of antimicrobial resistance of varying classes across the time series. Cumulative abundance (g) and overall richness (h) of antimicrobial resistance genes in patients did not differ between placebo and fecal transplant patients. We did observe possible increased abundance of (i) tetracycline and (j) aminoglycoside antibiotic resistance genes in placebo-treated patients compared with fecal transplant recipients.

**Fig S2.** Related to Figure 2: Fecal microbiota transplants in IBD patients alter community composition without affecting diversity. (a) Mean Bray-Curtis distance from the donor samples based on metagenomic species. Longitudinal changes in alpha diversity based on (b) Shannon diversity of metagenomic species, (c) ASV richness, (d) metagenomics species richness. (e) Bray-Curtis distance from baseline samples based on metagenomic species.

**Fig S3.** Related to Figure 3: longitudinal maintenance of transferred bacteria. (a) Tracking bacterial sources identified bacterial species transferred from donor to recipient as well as an invasion of bacteria of unknown origin. Each time-series plot indicates the fraction of total metagenomic species that were identified as from the patient or the donor, as shared between the two, or as from an unknown source. Regions of the graph colored as in Figure 1. (b) The counts of ASVs specific to the donor, specific to the patient at baseline, shared between the two, and categorized as transferred from the donor to the patient. This last category is a subset of the donor specific ASVs. (c) Persistent and temporary colonized bacteria based on metagenomic species.

**Fig S4.** Related to Figure 4: strain level transfer of donor bacteria. Using the flexible genome approach, we confirmed that samples from the same donor registered a strain match, while samples from different donors did not. Shown are the read depths of 1-kb windows of the reference genome for *Faecalibacterium prausnitzii* for (a) two samples from the same donor and (b) two samples from different donors, with red circles indicating genome segments that were present in one sample, but absent in the other. (c) Flexible genome strain matches against donor samples for *Bacteroides ovatus, B. caccae, B. fragilis, and P. merdae* indicated further instances of transferred donor strains in fecal transplant recipients. To evaluate our flexible genome method, we confirmed that (d) samples from placebo-treated patients did not match any of the donor samples and (e) samples from fecal transplant recipients who received an alternate donor did not match samples from that alternate donor except for isolated cases after the alternate donor period. StrainFinder analyses of (f) *B. fragilis*, (g) *B. thetaiotaomicron*, (h) *Bifidobacterium longum*, and (i) *Eubacterium eligens*.

**Fig S5.** Capsule delivery from an alternate donor introduced limited novel taxa. Tracking of ASVs as in Figure 3 revealed only limited that the transfer of unique taxa from the alternate donor via daily capsules. The majority of donor-transferred ASVs were shared between the two donors, but ASVs specific to the induction donor tended to outnumber those from the alternate donor.

**Fig S6.** Related to Figure 5: transfer of functional capacities. Plots of the frequencies of microbial functions according to source for patients FMT W, FMT P, and FMT F for genes related to (a) glycoside-hydrolases, (b) mucin digestion, (c) butyrate biosynthesis, (d) antimicrobial resistance, and (e) virulence factors. (f) Log10 cumulative abundance of all virulence factors in each patient across the trial period. (g) Cumulative abundance of virulence factors for all patients across the trial period. (h) Cumulative abundance of Proteobacteria in Patient V during the trial period.

**Fig S7.** Related to Figure 6: IgA coating of gut microbes. Some patients’ baseline samples had much greater correlation of IgA enrichment with a healthy donor than others’. Line plots (a) and corresponding Pearson *r* correlation values (b) for each patient at baseline. IgA enrichment did not correlate with a bacterium’s (c) abundance, (d) variance, or (e) bimodality. (f) IgA coating scores from each sample largely followed a normal distribution.

**Fig S8.** Phylogenetic tree of bacteria showing the IgA enrichment across samples and patients. Phylogenetic tree was constructed on the basis of 16S sequences. Each column is a heatmap corresponding to each patient, indicating the IgA coating score for that bacterium at a given time point.

**Table S1.** Metadata for stool samples collected, including patient clinical data and sequencing depths for 16S and shotgun metagenomic sequencing.

**Table S2.** Bacteria lost, gained, or maintained by placebo patients after the administration of antibiotics, according to 16S sequencing.

**Table S3.** Bacteria lost, gained, or maintained by placebo patients after the administration of antibiotics, according to metagenomic sequencing.

**Table S4.** Number of FMT patients receiving fecal material from Donor am (out of a total *n* = 4) who either (1) *shared* the ASV with the donor at baseline and after FMT, (2) showed *persistent* colonization of the ASV from the donor, or (3) showed *temporary* colonization.

**Table S5:** Transferred genes and associated taxa of butyrate biosynthesis genes.

**Table S6:** Additional metadata for each patient in the trial.

**Table S7:** Taxa identified as reliably IgA coated, IgA uncoated, and variably IgA coated or uncoated

## References and Notes

1. Z. Kassam, C. H. Lee, Y. Yuan, R. H. Hunt, Fecal microbiota transplantation for Clostridium difficile infection: systematic review and meta-analysis, Am. J. Gastroenterol. 108, 500–508 (2013).

2. E. van Nood, A. Vrieze, M. Nieuwdorp, S. Fuentes, E. G. Zoetendal, W. M. de Vos, C. E. Visser, E. J. Kuijper, J. F. W. M. Bartelsman, J. G. P. Tijssen, P. Speelman, M. G. W. Dijkgraaf, J. J. Keller, Duodenal Infusion of Donor Feces for Recurrent Clostridium difficile, N. Engl. J. Med. 368, 407–415 (2013).

3. L. M. Lix, L. A. Graff, J. R. Walker, I. Clara, P. Rawsthorne, L. Rogala, N. Miller, J. Ediger, T. Pretorius, C. N. Bernstein, Longitudinal study of quality of life and psychological functioning for active, fluctuating, and inactive disease patterns in inflammatory bowel disease, Inflamm. Bowel Dis. 14, 1575–1584 (2008).

4. X. C. Morgan, T. L. Tickle, H. Sokol, D. Gevers, K. L. Devaney, D. V. Ward, J. A. Reyes, S. A. Shah, N. LeLeiko, S. B. Snapper, A. Bousvaros, J. Korzenik, B. E. Sands, R. J. Xavier, C. Huttenhower, Dysfunction of the intestinal microbiome in inflammatory bowel disease and treatment, Genome Biol. 13, R79 (2012).

5. J. Halfvarson, C. J. Brislawn, R. Lamendella, Y. Vázquez-Baeza, W. A. Walters, L. M. Bramer, M. D’Amato, F. Bonfiglio, D. McDonald, A. Gonzalez, E. E. McClure, M. F. Dunklebarger, R. Knight, J. K. Jansson, Dynamics of the human gut microbiome in inflammatory bowel disease, Nat. Microbiol. 2, 17004 (2017).

6. S. Paramsothy, R. Paramsothy, D. T. Rubin, M. A. Kamm, N. O. Kaakoush, H. M. Mitchell, N. Castaño-Rodríguez, Faecal Microbiota Transplantation for Inflammatory Bowel Disease: A Systematic Review and Meta-analysis, J. Crohns Colitis 11, 1180–1199 (2017).

7. C. R. Kelly, S. Kahn, P. Kashyap, L. Laine, D. Rubin, A. Atreja, T. Moore, G. Wu, Update on Fecal Microbiota Transplantation 2015: Indications, Methodologies, Mechanisms, and Outlook, Gastroenterology 149, 223–237 (2015).

8. S. J. Ott, G. H. Waetzig, A. Rehman, J. Moltzau-Anderson, R. Bharti, J. A. Grasis, L. Cassidy, A. Tholey, H. Fickenscher, D. Seegert, P. Rosenstiel, S. Schreiber, Efficacy of Sterile Fecal Filtrate Transfer for Treating Patients With Clostridium difficile Infection, Gastroenterology 152, 799–811.e7 (2017).

9. D. R. Donohoe, N. Garge, X. Zhang, W. Sun, T. M. O’Connell, M. K. Bunger, S. J. Bultman, The Microbiome and Butyrate Regulate Energy Metabolism and Autophagy in the Mammalian Colon, Cell Metab. 13, 517–526 (2011).

10. M. S. Inan, R. J. Rasoulpour, L. Yin, A. K. Hubbard, D. W. Rosenberg, C. Giardina, The luminal short-chain fatty acid butyrate modulates NF-κB activity in a human colonic epithelial cell line, Gastroenterology 118, 724–734 (2000).

11. P. V. Chang, L. Hao, S. Offermanns, R. Medzhitov, The microbial metabolite butyrate regulates intestinal macrophage function via histone deacetylase inhibition, Proc. Natl. Acad. Sci. U. S. A. 111, 2247–2252 (2014).

12. Y. Furusawa, Y. Obata, S. Fukuda, T. A. Endo, G. Nakato, D. Takahashi, Y. Nakanishi, C. Uetake, K. Kato, T. Kato, M. Takahashi, N. N. Fukuda, S. Murakami, E. Miyauchi, S. Hino, K. Atarashi, S. Onawa, Y. Fujimura, T. Lockett, J. M. Clarke, D. L. Topping, M. Tomita, S. Hori, O. Ohara, T. Morita, H. Koseki, J. Kikuchi, K. Honda, K. Hase, H. Ohno, Commensal microbe-derived butyrate induces the differentiation of colonic regulatory T cells, Nature 504, 446–450 (2013).

13. M. D. Säemann, G. A. Böhmig, C. H. Österreicher, H. Burtscher, O. Parolini, C. Diakos, J. Stöckl, W. H. Hörl, G. J. Zlabinger, Anti-inflammatory effects of sodium butyrate on human monocytes: potent inhibition of IL-12 and up-regulation of IL-10 production, FASEB J. 14, 2380–2382 (2000).

14. N. Arpaia, C. Campbell, X. Fan, S. Dikiy, J. van der Veeken, P. deRoos, H. Liu, J. R. Cross, K. Pfeffer, P. J. Coffer, A. Y. Rudensky, Metabolites produced by commensal bacteria promote peripheral regulatory T-cell generation, Nature 504, 451–455 (2013).

15. B. Millan, H. Park, N. Hotte, O. Mathieu, P. Burguiere, T. A. Tompkins, D. Kao, K. L. Madsen, Fecal Microbial Transplants Reduce Antibiotic-resistant Genes in Patients With Recurrent Clostridium difficile Infection, Clin. Infect. Dis. 62, 1479–1486 (2016).

16. I. I. Ivanov, K. Atarashi, N. Manel, E. L. Brodie, T. Shima, U. Karaoz, D. Wei, K. C. Goldfarb, C. A. Santee, S. V. Lynch, T. Tanoue, A. Imaoka, K. Itoh, K. Takeda, Y. Umesaki, K. Honda, D. R. Littman, Induction of Intestinal Th17 Cells by Segmented Filamentous Bacteria, Cell 139, 485–498 (2009).

17. N. W. Palm, M. R. de Zoete, T. W. Cullen, N. A. Barry, J. Stefanowski, L. Hao, P. H. Degnan, J. Hu, I. Peter, W. Zhang, E. Ruggiero, J. H. Cho, A. L. Goodman, R. A. Flavell, Immunoglobulin A Coating Identifies Colitogenic Bacteria in Inflammatory Bowel Disease, Cell 158, 1000–1010 (2014).

18. M. G. de Agüero, S. C. Ganal-Vonarburg, T. Fuhrer, S. Rupp, Y. Uchimura, H. Li, A. Steinert, M. Heikenwalder, S. Hapfelmeier, U. Sauer, K. D. McCoy, A. J. Macpherson, The maternal microbiota drives early postnatal innate immune development, Science 351, 1296–1302 (2016).

19. S. Schwartz, I. Friedberg, I. V. Ivanov, L. A. Davidson, J. S. Goldsby, D. B. Dahl, D. Herman, M. Wang, S. M. Donovan, R. S. Chapkin, A metagenomic study of diet-dependent interaction between gut microbiota and host in infants reveals differences in immune response, Genome Biol. 13, r32 (2012).

20. P. Moayyedi, M. G. Surette, P. T. Kim, J. Libertucci, M. Wolfe, C. Onischi, D. Armstrong, J. K. Marshall, Z. Kassam, W. Reinisch, C. H. Lee, Fecal Microbiota Transplantation Induces Remission in Patients With Active Ulcerative Colitis in a Randomized Controlled Trial, Gastroenterology 149, 102–109.e6 (2015).

21. S. Paramsothy, M. A. Kamm, N. O. Kaakoush, A. J. Walsh, J. van den Bogaerde, D. Samuel, R. W. L. Leong, S. Connor, W. Ng, R. Paramsothy, W. Xuan, E. Lin, H. M. Mitchell, T. J. Borody, Multidonor intensive faecal microbiota transplantation for active ulcerative colitis: a randomised placebo-controlled trial, Lancet Lond. Engl. 389, 1218–1228 (2017).

22. N. G. Rossen, S. Fuentes, M. J. van der Spek, J. G. Tijssen, J. H. A. Hartman, A. Duflou, M. Löwenberg, G. R. van den Brink, E. M. H. Mathus-Vliegen, W. M. de Vos, E. G. Zoetendal, G. R. D’Haens, C. Y. Ponsioen, Findings From a Randomized Controlled Trial of Fecal Transplantation for Patients With Ulcerative Colitis, Gastroenterology 149, 110–118.e4 (2015).

23. S. P. Costello, P. A. Hughes, O. Waters, R. V. Bryant, A. D. Vincent, P. Blatchford, R. Katsikeros, J. Makanyanga, M. A. Campaniello, C. Mavrangelos, C. P. Rosewarne, C. Bickley, C. Peters, M. N. Schoeman, M. A. Conlon, I. C. Roberts-Thomson, J. M. Andrews, Effect of Fecal Microbiota Transplantation on 8-Week Remission in Patients With Ulcerative Colitis: A Randomized Clinical Trial, JAMA 321, 156–164 (2019).

24. H. Sokol, B. Pigneur, L. Watterlot, O. Lakhdari, L. G. Bermúdez-Humarán, J.-J. Gratadoux, S. Blugeon, C. Bridonneau, J.-P. Furet, G. Corthier, C. Grangette, N. Vasquez, P. Pochart, G. Trugnan, G. Thomas, H. M. Blottière, J. Doré, P. Marteau, P. Seksik, P. Langella, Faecalibacterium prausnitzii is an anti-inflammatory commensal bacterium identified by gut microbiota analysis of Crohn disease patients, Proc. Natl. Acad. Sci. 105, 16731–16736 (2008).

25. E. Bolyen, J. R. Rideout, M. R. Dillon, N. A. Bokulich, C. Abnet, G. A. Al-Ghalith, H. Alexander, E. J. Alm, M. Arumugam, F. Asnicar, Y. Bai, J. E. Bisanz, K. Bittinger, A. Brejnrod, C. J. Brislawn, C. T. Brown, B. J. Callahan, A. M. Caraballo-Rodríguez, J. Chase, E. Cope, R. D. Silva, P. C. Dorrestein, G. M. Douglas, D. M. Durall, C. Duvallet, C. F. Edwardson, M. Ernst, M. Estaki, J. Fouquier, J. M. Gauglitz, D. L. Gibson, A. Gonzalez, K. Gorlick, J. Guo, B. Hillmann, S. Holmes, H. Holste, C. Huttenhower, G. Huttley, S. Janssen, A. K. Jarmusch, L. Jiang, B. Kaehler, K. B. Kang, C. R. Keefe, P. Keim, S. T. Kelley, D. Knights, I. Koester, T. Kosciolek, J. Kreps, M. G. Langille, J. Lee, R. Ley, Y.-X. Liu, E. Loftfield, C. Lozupone, M. Maher, C. Marotz, B. D. Martin, D. McDonald, L. J. McIver, A. V. Melnik, J. L. Metcalf, S. C. Morgan, J. Morton, A. T. Naimey, J. A. Navas-Molina, L. F. Nothias, S. B. Orchanian, T. Pearson, S. L. Peoples, D. Petras, M. L. Preuss, E. Pruesse, L. B. Rasmussen, A. Rivers, I. I. Michael S Robeson, P. Rosenthal, N. Segata, M. Shaffer, A. Shiffer, R. Sinha, S. J. Song, J. R. Spear, A. D. Swafford, L. R. Thompson, P. J. Torres, P. Trinh, A. Tripathi, P. J. Turnbaugh, S. Ul-Hasan, J. J. van der Hooft, F. Vargas, Y. Vázquez-Baeza, E. Vogtmann, M. von Hippel, W. Walters, Y. Wan, M. Wang, J. Warren, K. C. Weber, C. H. Williamson, A. D. Willis, Z. Z. Xu, J. R. Zaneveld, Y. Zhang, Q. Zhu, R. Knight, J. G. Caporaso, QIIME 2: Reproducible, interactive, scalable, and extensible microbiome data science (PeerJ Inc., 2018; https://peerj.com/preprints/27295).

26. B. J. Callahan, P. J. McMurdie, M. J. Rosen, A. W. Han, A. J. A. Johnson, S. P. Holmes, DADA2: High-resolution sample inference from Illumina amplicon data, Nat. Methods 13, 581–583 (2016).

27. D. T. Truong, E. A. Franzosa, T. L. Tickle, M. Scholz, G. Weingart, E. Pasolli, A. Tett, C. Huttenhower, N. Segata, MetaPhlAn2 for enhanced metagenomic taxonomic profiling, Nat. Methods 12, 902–903 (2015).

28. A. Palleja, K. H. Mikkelsen, S. K. Forslund, A. Kashani, K. H. Allin, T. Nielsen, T. H. Hansen, S. Liang, Q. Feng, C. Zhang, P. T. Pyl, L. P. Coelho, H. Yang, J. Wang, A. Typas, M. F. Nielsen, H. B. Nielsen, P. Bork, J. Wang, T. Vilsbøll, T. Hansen, F. K. Knop, M. Arumugam, O. Pedersen, Recovery of gut microbiota of healthy adults following antibiotic exposure, Nat. Microbiol. 3, 1255 (2018).

29. C. Staley, C. R. Kelly, L. J. Brandt, A. Khoruts, M. J. Sadowsky, Complete Microbiota Engraftment Is Not Essential for Recovery from Recurrent Clostridium difficile Infection following Fecal Microbiota Transplantation, mBio 7, e01965–16 (2016).

30. C. R. Kelly, A. Khoruts, C. Staley, M. J. Sadowsky, M. Abd, M. Alani, B. Bakow, P. Curran, J. McKenney, A. Tisch, S. E. Reinert, J. T. Machan, L. J. Brandt, Effect of Fecal Microbiota Transplantation on Recurrence in Multiply Recurrent Clostridium difficile Infection: A Randomized Trial, Ann. Intern. Med. 165, 609–616 (2016).

31. C. S. Smillie, J. Sauk, D. Gevers, J. Friedman, J. Sung, I. Youngster, E. L. Hohmann, C. Staley, A. Khoruts, M. J. Sadowsky, J. R. Allegretti, M. B. Smith, R. J. Xavier, E. J. Alm, Strain Tracking Reveals the Determinants of Bacterial Engraftment in the Human Gut Following Fecal Microbiota Transplantation, Cell Host Microbe 23, 229–240.e5 (2018).

32. S. S. Li, A. Zhu, V. Benes, P. I. Costea, R. Hercog, F. Hildebrand, J. Huerta-Cepas, M. Nieuwdorp, J. Salojärvi, A. Y. Voigt, G. Zeller, S. Sunagawa, W. M. de Vos, P. Bork, Durable coexistence of donor and recipient strains after fecal microbiota transplantation, Science 352, 586–589 (2016).

33. G. Guyatt, A. Mitchell, E. J. Irvine, J. Singer, N. Williams, R. Goodacre, C. Tompkins, A new measure of health status for clinical trials in inflammatory bowel disease, Gastroenterology 96, 804–810 (1989).

34. I. L. Brito, T. Gurry, S. Zhao, K. Huang, S. K. Young, T. P. Shea, W. Naisilisili, A. P. Jenkins, S. D. Jupiter, D. Gevers, E. J. Alm, Transmission of human-associated microbiota along family and social networks, Nat. Microbiol., 1 (2019).

35. C. Duvallet, S. M. Gibbons, T. Gurry, R. A. Irizarry, E. J. Alm, Meta-analysis of gut microbiome studies identifies disease-specific and shared responses, Nat. Commun. 8, 1784 (2017).

36. E. R. Dubberke, K. M. Mullane, D. N. Gerding, C. H. Lee, T. J. Louie, H. Guthertz, C. Jones, Clearance of Vancomycin-Resistant Enterococcus Concomitant With Administration of a Microbiota-Based Drug Targeted at Recurrent Clostridium difficile Infection, Open Forum Infect. Dis. 3 (2016), doi:10.1093/ofid/ofw133.

37. Y. Wei, J. Gong, W. Zhu, D. Guo, L. Gu, N. Li, J. Li, Fecal microbiota transplantation restores dysbiosis in patients with methicillin resistant Staphylococcus aureus enterocolitis, BMC Infect. Dis. 15, 265 (2015).

38. O. Pabst, New concepts in the generation and functions of IgA, Nat. Rev. Immunol. 12, 821–832 (2012).

39. G. P. Donaldson, M. S. Ladinsky, K. B. Yu, J. G. Sanders, B. B. Yoo, W.-C. Chou, M. E. Conner, A. M. Earl, R. Knight, P. J. Bjorkman, S. K. Mazmanian, Gut microbiota utilize immunoglobulin A for mucosal colonization, Science 360, 795–800 (2018).

40. K. McLoughlin, J. Schluter, S. Rakoff-Nahoum, A. L. Smith, K. R. Foster, Host Selection of Microbiota via Differential Adhesion, Cell Host Microbe 19, 550–559 (2016).

41. D. A. Peterson, N. P. McNulty, J. L. Guruge, J. I. Gordon, IgA Response to Symbiotic Bacteria as a Mediator of Gut Homeostasis, Cell Host Microbe 2, 328–339 (2007).

42. S. M. Kearney, S. M. Gibbons, S. E. Erdman, E. J. Alm, Orthogonal Dietary Niche Enables Reversible Engraftment of a Gut Bacterial Commensal, Cell Rep. 24, 1842–1851 (2018).

43. A. B. Hall, M. Yassour, J. Sauk, A. Garner, X. Jiang, T. Arthur, G. K. Lagoudas, T. Vatanen, N. Fornelos, R. Wilson, M. Bertha, M. Cohen, J. Garber, H. Khalili, D. Gevers, A. N. Ananthakrishnan, S. Kugathasan, E. S. Lander, P. Blainey, H. Vlamakis, R. J. Xavier, C. Huttenhower, A novel Ruminococcus gnavus clade enriched in inflammatory bowel disease patients, Genome Med. 9, 103 (2017).

44. M. M. Wegorzewska, R. W. P. Glowacki, S. A. Hsieh, D. L. Donermeyer, C. A. Hickey, S. C. Horvath, E. C. Martens, T. S. Stappenbeck, P. M. Allen, Diet modulates colonic T cell responses by regulating the expression of a Bacteroides thetaiotaomicron antigen, Sci. Immunol. 4, eaau9079 (2019).

45. C. Prantera, F. Zannoni, M. L. Scribano, E. Berto, A. Andreoli, A. Kohn, C. Luzi, An antibiotic regimen for the treatment of active Crohn’s disease: a randomized, controlled clinical trial of metronidazole plus ciprofloxacin, Am. J. Gastroenterol. 91, 328–332 (1996).

46. K. J. Khan, T. A. Ullman, A. C. Ford, M. T. Abreu, A. Abadir, A. Abadir, J. K. Marshall, N. J. Talley, P. Moayyedi, Antibiotic therapy in inflammatory bowel disease: a systematic review and meta-analysis, Am. J. Gastroenterol. 106, 661–673 (2011).

47. S. Ben-Horin, M. Margalit, P. Bossuyt, J. Maul, Y. Shapira, D. Bojic, I. Chermesh, A. Al-Rifai, A. Schoepfer, M. Bosani, M. Allez, P. L. Lakatos, F. Bossa, A. Eser, T. Stefanelli, F. Carbonnel, K. Katsanos, D. Checchin, I. S. de Miera, Y. Chowers, G. W. Moran, Combination Immunomodulator and Antibiotic Treatment in Patients With Inflammatory Bowel Disease and Clostridium difficile Infection, Clin. Gastroenterol. Hepatol. 7, 981–987 (2009).

48. A. H. Steinhart, B. G. Feagan, C. J. Wong, M. Vandervoort, S. Mikolainis, K. Croitoru, E. Seidman, D. J. Leddin, A. Bitton, E. Drouin, A. Cohen, G. R. Greenberg, Combined budesonide and antibiotic therapy for active Crohn’s disease: A randomized controlled trial, Gastroenterology 123, 33–40 (2002).

49. C. Prantera, H. Lochs, M. Campieri, M. L. Scribano, G. C. Sturniolo, F. Castiglione, M. Cottone, Antibiotic treatment of Crohn’s disease: results of a multicentre, double blind, randomized, placebo-controlled trial with rifaximin, Aliment. Pharmacol. Ther. 23, 1117–1125 (2006).

50. T. Ohkusa, K. Kato, S. Terao, T. Chiba, K. Mabe, K. Murakami, Y. Mizokami, T. Sugiyama, A. Yanaka, Y. Takeuchi, S. Yamato, T. Yokoyama, I. Okayasu, S. Watanabe, H. Tajiri, N. Sato, Japan UC Antibiotic Therapy Study Group, Newly developed antibiotic combination therapy for ulcerative colitis: a double-blind placebo-controlled multicenter trial, Am. J. Gastroenterol. 105, 1820–1829 (2010).

51. E. S. Shepherd, W. C. DeLoache, K. M. Pruss, W. R. Whitaker, J. L. Sonnenburg, An exclusive metabolic niche enables strain engraftment in the gut microbiota, Nature 557, 434–438 (2018).

52. E. D. Sonnenburg, H. Zheng, P. Joglekar, S. K. Higginbottom, S. J. Firbank, D. N. Bolam, J. L. Sonnenburg, Specificity of polysaccharide use in intestinal bacteroides species determines diet-induced microbiota alterations, Cell 141, 1241–1252 (2010).

53. S. Wang, M. Xu, W. Wang, X. Cao, M. Piao, S. Khan, F. Yan, H. Cao, B. Wang, Systematic Review: Adverse Events of Fecal Microbiota Transplantation, PLOS ONE 11, e0161174 (2016).

54. C. S. Smillie, M. B. Smith, J. Friedman, O. X. Cordero, L. A. David, E. J. Alm, Ecology drives a global network of gene exchange connecting the human microbiome, Nature 480, 241–244 (2011).

55. E. W. Rogier, A. L. Frantz, M. E. C. Bruno, C. S. Kaetzel, Secretory IgA is Concentrated in the Outer Layer of Colonic Mucus along with Gut Bacteria, Pathogens 3, 390–403 (2014).

56. J. J. Bunker, S. A. Erickson, T. M. Flynn, C. Henry, J. C. Koval, M. Meisel, B. Jabri, D. A. Antonopoulos, P. C. Wilson, A. Bendelac, Natural polyreactive IgA antibodies coat the intestinal microbiota, Science 358, eaan6619 (2017).

57. J. G. Caporaso, C. L. Lauber, W. A. Walters, D. Berg-Lyons, C. A. Lozupone, P. J. Turnbaugh, N. Fierer, R. Knight, Global patterns of 16S rRNA diversity at a depth of millions of sequences per sample, Proc. Natl. Acad. Sci. 108, 4516–4522 (2011).

58. C. Quast, E. Pruesse, P. Yilmaz, J. Gerken, T. Schweer, P. Yarza, J. Peplies, F. O. Glöckner, The SILVA ribosomal RNA gene database project: improved data processing and web-based tools, Nucleic Acids Res. 41, D590–D596 (2013).

59. J. Kaminski, M. K. Gibson, E. A. Franzosa, N. Segata, G. Dantas, C. Huttenhower, High-Specificity Targeted Functional Profiling in Microbial Communities with ShortBRED, PLOS Comput. Biol. 11, e1004557 (2015).

60. M. Vital, A. Karch, D. H. Pieper, Colonic Butyrate-Producing Communities in Humans: an Overview Using Omics Data, mSystems 2 (2017), doi:10.1128/mSystems.00130-17.

61. D. A. Ravcheev, I. Thiele, Comparative Genomic Analysis of the Human Gut Microbiome Reveals a Broad Distribution of Metabolic Pathways for the Degradation of Host-Synthetized Mucin Glycans and Utilization of Mucin-Derived Monosaccharides, Front. Genet. 8 (2017), doi:10.3389/fgene.2017.00111.

62. V. Lombard, H. Golaconda Ramulu, E. Drula, P. M. Coutinho, B. Henrissat, The carbohydrate-active enzymes database (CAZy) in 2013, Nucleic Acids Res. 42, D490–D495 (2014).

63. B. Jia, A. R. Raphenya, B. Alcock, N. Waglechner, P. Guo, K. K. Tsang, B. A. Lago, B. M. Dave, S. Pereira, A. N. Sharma, S. Doshi, M. Courtot, R. Lo, L. E. Williams, J. G. Frye, T. Elsayegh, D. Sardar, E. L. Westman, A. C. Pawlowski, T. A. Johnson, F. S. L. Brinkman, G. D. Wright, A. G. McArthur, CARD 2017: expansion and model-centric curation of the comprehensive antibiotic resistance database, Nucleic Acids Res. 45, D566–D573 (2017).

64. L. Chen, D. Zheng, B. Liu, J. Yang, Q. Jin, VFDB 2016: hierarchical and refined dataset for big data analysis—10 years on, Nucleic Acids Res. 44, D694–D697 (2016).

65. H. Li, R. Durbin, Fast and accurate long-read alignment with Burrows-Wheeler transform, Bioinforma. Oxf. Engl. 26, 589–595 (2010).

66. M. Wu, J. A. Eisen, A simple, fast, and accurate method of phylogenomic inference, Genome Biol. 9, R151 (2008).

67. H. Li, B. Handsaker, A. Wysoker, T. Fennell, J. Ruan, N. Homer, G. Marth, G. Abecasis, R. Durbin, 1000 Genome Project Data Processing Subgroup, The Sequence Alignment/Map format and SAMtools, Bioinforma. Oxf. Engl. 25, 2078–2079 (2009).

68. A. Amir, D. McDonald, J. A. Navas-Molina, E. Kopylova, J. T. Morton, Z. Z. Xu, E. P. Kightley, L. R. Thompson, E. R. Hyde, A. Gonzalez, R. Knight, Deblur Rapidly Resolves Single-Nucleotide Community Sequence Patterns, mSystems 2, e00191–16 (2017).

69. M. N. Price, P. S. Dehal, A. P. Arkin, FastTree 2 – Approximately Maximum-Likelihood Trees for Large Alignments, PLOS ONE 5, e9490 (2010).

70. I. Letunic, P. Bork, Interactive Tree Of Life (iTOL) v4: recent updates and new developments, Nucleic Acids Res., doi:10.1093/nar/gkz239.

